# Reconstitution of the Spinal Cord Injury Microenvironment in Adult Neural Stem Cell-Derived Organoids

**DOI:** 10.64898/2026.03.14.711818

**Authors:** Martyna Lukoseviciute, Vlada-Iuliana Panfil, Timm Häneke, Anaïs Julien, Enric Llorens-Bobadilla, Christian Göritz, Jonas Frisén

## Abstract

Following spinal cord injury, endogenous neural stem cells (NSCs) derived from ependymal cells become activated but fail to functionally regenerate the tissue, largely because the injury microenvironment constrains their differentiation toward glial fates. Dissecting how specific niche components drive these outcomes has remained challenging in vivo, and current neural organoid models predominantly recapitulate embryonic neurodevelopment rather than the adult injury context. Here we describe neuroids - a modular organoid system built from injury-activated adult spinal cord ependymal NSCs that spontaneously differentiate into neurons, astrocytes, and to some degree oligodendrocytes within a self-organised 3D structure. Using a bottom-up approach, we reconstruct the injury niche by incorporating meningeal fibroblasts and primary adult microglia, individually and in combination. Fibroblasts accumulate in the organoid core, deposit extracellular matrix (ECM), and trigger reactive astrocyte responses mirroring in vivo scar organisation, while microglia integrate throughout, adopt heterogeneous activation states, and remain functionally active. Their combined incorporation further enhances ECM deposition and promotes oligodendrocyte lineage commitment, suggesting cooperative niche interactions. Single-nucleus multiome profiling and trajectory inference show that these injury-like conditions shift NSC differentiation away from neuronal programs toward proliferative and astroglial states, recapitulating NSC behaviour after injury in vivo. Ligand-receptor analysis implicates microglia-derived TGFβ, WNT, and ECM-associated signals as candidate drivers of this gliogenic bias. Together, neuroids provide a tractable platform to study how the adult injury niche regulates endogenous NSC fate, and to identify strategies that simultaneously redirect these cells toward regeneration while targeting the fibrotic scar - two barriers that together prevent functional recovery after spinal cord injury.

## Introduction

Traumatic spinal cord injury (SCI) leads to irreversible neurological deficits because the adult mammalian central nervous system has limited regenerative capacity. Following injury, the lesion site rapidly becomes populated by reactive astrocytes, activated immune cells, and fibroblasts and pericytes that together generate a dense inflammatory and extracellular matrix-rich profibrotic environment. While these responses stabilise damaged tissue and limit secondary damage, they also create molecular and physical barriers that restrict neuronal regeneration and axonal growth^1–3^.

Despite this limited regenerative outcome, the adult spinal cord was found to harbour endogenous neural stem cells (eNSCs) derived from ependymal cells lining the central canal. These cells become activated following injury, proliferate, and generate multiple neural lineages^4–6^. Lineage tracing studies have demonstrated that ependymal-derived NSCs can differentiate into astrocytes, seldomly oligodendrocytes and, in neurogenic niches or in vitro, neurons^4,7–10^. However, in vivo spinal cord their differentiation is strongly biased toward astrocytic scar formation rather than neuronal regeneration, which while contributes to prevention of secondary injury spreading, does not lead to a functional tissue regeneration^11^. These observations suggest that while ependymal cells retain intrinsic multilineage potential, their cell fate decisions are constrained by the surrounding spinal cord injury microenvironment.

Hence, learning how to modulate endogenous neural stem cell responses, for instance via modifying their niche or intrinsic gene regulation holds a promise for regenerative SCI strategies. Cell transplantation approaches using neural stem or progenitor cells have shown promise in experimental models, but their clinical translation remains challenging due to limited graft survival, immune rejection, and the persistence of a hostile injury microenvironment that can restrict neuronal differentiation or integration into host circuits ^12–14^. Therefore, experimental strategies that instead harness endogenous stem cells may offer advantages, because these cells are already integrated within the tissue architecture, become transiently activated following injury and do not lead to immune rejection. However, it remains critical to learn how to manipulate eNSC fates in the injury environment, for examples by promoting neurogenesis, and the secret in doing so might be found in their microenvironment.

Several components of the injury niche have already been identified in shaping neural stem cell behaviour. For instance, activated microglia and other infiltrating immune cells release cytokines that might activate neural stem cell properties, but suppress neurogenesis or reinforce gliogenic differentiation programs^15–20^. In parallel, meningeal fibroblasts and pericytes invade the lesion core and deposit extracellular matrix (ECM) proteins that form a fibrotic scar, which contributes to lesion stabilisation but also serves as limiting permanent barrier to axon regeneration or new neuronal cell integration^3,21,22^. Despite these experimental observations, it remains difficult to dissect how individual environmental signals influence endogenous neural stem cell behaviour in vivo due to the complex spatiotemporal cellular heterogeneity within the injury environment.

Three-dimensional organoid systems have emerged as powerful tools to model cellular interactions in controlled environments. Most neural organoids developed to date recapitulate aspects of embryonic neurodevelopment and have provided insight into neural patterning, lineage specification, and disease mechanisms^23–27^. More recently, organoid systems have been improved further to recapitulate wholistic tissue level complexity by incorporating immune or stromal components^28–30^. However, organoid models specifically designed to investigate the adult spinal cord injury niche remain limited.

Here we establish an adult spinal cord neural stem cell-derived organoid system, which we term a neuroid, to study how environmental cues regulate neural stem cell behaviour. In contrast to developmental organoids, this system is derived from injury-activated adult spinal cord ependymal stem cells and therefore models endogenous regenerative cell populations.

Using a bottom-up approach, we reconstruct key components of the spinal cord injury niche, including fibrotic scar-forming fibroblasts and inflammatory microglia. We use this system to also examine how the environmental signals influence neural stem cell differentiation trajectories. Through single-nuclei multiome profiling and lineage trajectory analysis we show that reconstructed injury-like environments shift neural stem cell differentiation away from neuronal programs and toward proliferative and astroglial states. These findings establish a tractable platform to dissect how environmental signals regulate endogenous neural stem cell responses following spinal cord injury.

## Materials and Methods

### Animals and spinal cord injury

All experimental procedures involving male and female wild-type C57BL/6 mice were conducted in strict compliance with ethical guidelines approved by the Stockholms Norra Djurförsöksetiska Nämnd (Karolinska Institute, Stockholm, Sweden). Mice were housed in individually ventilated cages (2–5 per cage) under a 12-hour light/dark cycle with ad libitum access to food and water. Surgical spinal cord injury was induced via a dorsal funiculus incision under isoflurane anesthesia (4% induction, 2% maintenance). Pre-emptive analgesia was administered subcutaneously, consisting of buprenorphine (0.1 mg/kg; Schering-Plough) and carprofen (5 mg/kg; Pfizer). Following a laminectomy at the T9 vertebral level, a transverse incision was made across the dorsal funiculus and extended rostrally to span one spinal segment using microsurgical scissors. Post-operatively, animals were monitored until the return of spontaneous breathing and placed in pre-warmed cages for recovery.

### Neural stem cell isolation and neurosphere generation

Neural stem cells were isolated from the spinal cords of male and female C57BL/6 wild-type mice two days following a dorsal funiculus incision. Immediately after dissection, tissues were maintained in ice-cold DPBS (Gibco, Cat. No. 14287080) until processing. Tissue dissociation into single-cell suspensions was performed using the gentleMACS Neural Dissociation Kit (Miltenyi Biotec, Cat. No. 130-092-628), utilising papain-based enzymatic digestion according to the manufacturer’s protocol. To ensure high purity, myelin debris was removed using Myelin Removal Beads II (Miltenyi Biotec, Cat. No. 130-096-433).

Isolated cells were cultured in proliferation medium consisting of DMEM/F12 (Thermo Fisher Scientific, Cat. No. 10567014, 31765035) supplemented with 2% B27 (Thermo Fisher Scientific, Cat. No. 17504044), 1% Penicillin/Streptomycin (Thermo Fisher Scientific, Cat. No. 15140148), 1 μg/mL Fungizone (Thermo Fisher Scientific, Cat. No. 15290018), 20 ng/mL epidermal growth factor (EGF; Sigma-Aldrich, Cat. No. E9644), and 25 ng/mL fibroblast growth factor 2 (FGF2; Sigma-Aldrich, Cat. No. F0291). Cells were seeded in 24-well plates and incubated at 37°C with 5% CO_2_. To sustain proliferation, growth factors were replenished on days 4, 7, and 10, with additional B27 supplement added on day 7. Neurospheres were cultured for 7–10 days before passaging at least once before neuroid generation using Accutase containing EDTA (STEMCELL Technologies, Cat. No. 07920).

### Neuroid generation

Neurospheres were enzymatically dissociated into single-cell suspensions using Accutase and filtered through 70-μm cell strainers to remove residual aggregates. The resulting cells were resuspended in proliferation medium supplemented with 20% sterile-filtered conditioned media (filtered through 20-μm syringe filters). Cells were counted and seeded at a density of 5,000 cells per well in 96-well ultra-low attachment plates (Thermo Fisher Scientific, Cat. No. 174925) in 250-μL of proliferation medium (as described above) per well. To facilitate uniform aggregation, plates were centrifuged at 100 × g for 4 minutes immediately after seeding. Cultures were incubated at 37°C in 5% CO_2_ for 7 days, with a 125-μL partial medium exchange performed on day 4 to maintain nutrient levels.

### Fibroblast isolation and co-culture

Fibroblasts were isolated from Col1a1-CreERT; tdTomato dual-reporter transgenic mice, a model that enables tamoxifen-inducible Cre recombinase expression in collagen I-positive cells, thereby activating the fluorescent tdTomato reporter. To induce Cre recombination, tamoxifen powder (Sigma-Aldrich, Cat. No. T5648) was prepared in corn oil at 20mg/ml and dissolved at 60°C for 2 hours. Mice were injected intraperitoneally with 2mg of tamoxifen for 3 consecutive days. 2-3 mice were euthanized by pentobarbital overdose, and brains were dissected on ice-cold PBS. After removing the skin, the skull was dissected on ice and meninges were collected using Dumont no. 5 forceps (Merck, Cat. No. F6521-1EA) and directly transferred into digesting medium containing DMEM (Life Technologies, Cat. No.61965-026) with 1% Collagenase D (Roche, Cat. No. 11088866001) and 1% trypsin (Life Technologies, Cat. No. 15090-046,). Tissue was digested for 1h at 37°C on a rotator in 2 ml of digesting medium. Every 20 min, cell suspension was pipetted up and down to aid cell dissociation. Digestion was stopped by adding 10 ml of growth media containing DMEM with 10% fetal bovine serum (Life Technologies, Cat. No. A5256701) and 1% Penicillin/Streptomycin. Cells were then filtered through 40 μm filters (Dutscher, Cat. No. 352340), centrifuged 10 min at 300×g and resuspended in 3 mL of growth medium prior plating in 6-well plates (Starstedt, Cat. No. 83.3920). Medium was changed 24 h after plating, and every 2-3 days before cells reach 80% confluence. Cells were passaged between 1 and 3 rounds prior sorting. For cell sorting, cells were resuspended in 1 mL of growth medium. Cell viability marker Sytox blue (1:1000 dilution, Thermofischer, Cat. No. S34857) was added 5 min before sorting. Col1a1-derived tdTomato+ cells were sorting using BD FACS Aria Fusion (BD Biosciences) and collected in neuroid differentiation medium and co-cultured with neuroids at a density of 500–1,000 cells per organoid.

To specifically capture fibroblasts with active collagen I synthesis, we additionally incorporated a Col1a1-GFP reporter line, allowing for the sorting of Col1a1-GFP+ cells same way as described above for neuroid co-cultures.

### Microglia isolation, expansion, and co-culture

Microglia were isolated and expanded using a modified protocol^31^. Briefly, adult brains were dissected and dissociated using the Adult Brain Dissociation Kit (Miltenyi Biotec, Cat. No. 130-107-677) according to the manufacturer’s instructions. The resulting cell suspension was resuspended in 7.5 mL of microglia expansion medium consisting of DMEM supplemented with 10% fetal bovine serum (FBS), 1% Penicillin/Streptomycin, 100 ng/mL macrophage colony-stimulating factor (M-CSF; Thermo Fisher Scientific, Cat. No. 315-02-50UG), 50 ng/mL granulocyte-macrophage colony-stimulating factor (GM-CSF; Thermo Fisher Scientific, Cat. No. 315-03-20UG), and 25% sterile fibroblast-conditioned medium to specifically enhance microglia cell yield. Cells were seeded in T75 flasks (Greiner, Cat. No. C7231) and maintained with a half-volume medium exchange on day 5 and a full media exchange on day 10. On day 12, cultures were topped up to 10 mL and agitated on an orbital shaker at 180 rpm for 2.5 hours at 37°C (0% CO_2_) to selectively detach microglia from the underlying astrocyte monolayer. The supernatant containing detached microglia was collected and centrifuged at 300 × g for 5 minutes.

Prior to neuroid co-culture, microglial identity and purity were validated via immunofluorescent staining for Iba1, CD45, and Ki67, and quantified by flow cytometry (Cd45-Brilliant Violet 650, BioLegend, Cat. No. 109836; Cd11b-PE-Cyanine7, Invitrogen, Cat. No. A15848). For surface staining, detached cells were incubated with antibody cocktails in 96-well plates for 20 minutes on ice, alongside unstained, single-stained, and fluorescence-minus-one (FMO) controls. Following a PBS wash, cells were analysed on BD Fusion Aria III using a hierarchical gating strategy: populations were isolated by forward/side scatter (FSC-A/SSC-A), singlets were selected (FSC-H/FSC-A), and viability was confirmed using live-dead stain Zombie Aqua (1:1000 dilution, BioLegend, Cat. No. 423101). Microglia were defined as live (Zombie Aqua -), CD45+/CD11b+ double-positive cells. Microglia were then co-cultured with neuroids at 1,000 cells per organoid; remaining cells were re-plated for subsequent expansion in culture as described above. To minimise animal use, microglia and neural stem cells were isolated in parallel from the brain and spinal cord of the same animal, respectively.

For live-cell imaging experiments, microglia were labelled with CellTracker™ Red CMTPX (Invitrogen, Cat. No. C34552). Cells were incubated in serum-free medium containing 10 μM dye for 30–50 minutes at 37°C, pelleted at 300 × g for 10 minutes, and resuspended in differentiation medium. Labelled microglia were co-seeded with neuroids and monitored using the IncuCyte® live-cell analysis system (Sartorius) to track microglial migration and interactions with the neuroid surface over time.

### Neuroid differentiation and microenvironment reconstruction

Neural differentiation was induced by replacing proliferation medium with neuroid differentiation medium, composed of Neurobasal medium (Thermo Fisher Scientific, Cat. No. A3582901) supplemented with 1% B27, 1% Penicillin/Streptomycin, 0.5 μg/mL Fungizone, 1% GlutaMAX (Cat. No. 35050061), 0.5% N2 (Cat. No. 17502048), and 0.5% MEM-NEAA (Cat. No. 11140050). Medium exchanges (125 μL) were performed every other day throughout the differentiation period.

To reconstruct specific microenvironments, distinct co-culture protocols were established. For fibrotic neuroids, 500–1,000 sorted fibroblasts were added per organoid on day 2 of differentiation, following an initial 7-day proliferation phase. For immune-neuroids, 1,000 microglia were seeded per organoid on day 2 of differentiation using the same basal protocol. To model a fibro-inflammatory niche, a sequential addition strategy was employed: microglia were introduced on day 2 of differentiation, followed by the addition of 500–1,000 fibroblasts on day 4. Organoid morphology and cellular integration were monitored via bright-field microscopy using 5× and 10× air objectives on a Zeiss Axio Observer system.

### Organoid whole-mount staining and imaging

Organoids were harvested in PCR tubes, washed twice with PBS, and fixed in 4% formaldehyde (in PBS, SigmaAldrich, Cat. No. 423101) for 1 hour at room temperature (RT). Following two PBS washes, samples were incubated in 0.1 M glycine (in PBS, Sigma Aldrich, Cat.No. G8790) for 1 hour at RT to quench autofluorescence. For permeabilization and clearing, organoids were incubated in 4% sodium dodecyl sulfate (SDS; Thermo Fisher Scientific, Cat. No. AM9820) in PBS for 4 hours at 50°C with agitation (300 rpm), followed by a 1-hour PBS wash. Immunostaining was performed in PBS containing 0.1% Tween-20 (PBST, Sigma Aldrich, Cat. No. P6585). Samples were incubated with primary antibodies for 24 hours at RT, washed for 24 hours, and then incubated with secondary antibodies and DAPI. Prior to mounting, organoids were optically cleared in Omnipaque™ (GE Healthcare, NDC 0407-1414-89) for 12 hours. Images were acquired using a Zeiss LSM700 confocal microscope with 10× and 20× air objectives. A complete list of antibodies is provided in Supplementary Table 1.

### Organoid cryosection staining and imaging

For cryosectioning, organoids were fixed as described above, embedded in OCT compound (HistoLab, Cat. No. 45830) and sectioned at a thickness of 20 μm. Sections were washed twice with PBST (0.2% Tween-20 in PBS) for 5 minutes and blocked for 2 hours in blocking buffer (5% goat serum, 0.2% Triton X-100 in PBS). Primary antibodies were incubated in blocking buffer for 24 hours at 4°C. Sections were then washed three times (5 minutes each) in PBS, incubated with secondary antibodies for 1 hour at RT, and washed again three times. Nuclear counterstaining was performed with DAPI for 5 minutes, followed by two final PBS washes before mounting. Images were acquired using a Zeiss LSM700 confocal microscope with 10× and 20× air objectives. A complete list of antibodies is provided in Supplementary Table 1.

### EdU cell proliferation assay

To assess cellular proliferation within neuroids, the Click-iT™ EdU Cell Proliferation Kit for Imaging (Invitrogen, Cat. No. C10340) was employed according to the manufacturer’s instructions. Briefly, 5-ethynyl-2′-deoxyuridine (EdU) was added directly to the organoid culture medium at a final volume of 20 μL per well. Organoids were incubated with EdU for 24 to 48 hours, depending on the specific experimental timeline. Following incubation, samples were fixed in 4% formaldehyde as described in the whole-mount staining protocol. The Click-iT reaction cocktail containing Alexa Fluor™ 647 azide was applied to label incorporated EdU. Subsequently, samples underwent standard whole-mount immunostaining for cell-type-specific markers and nuclear counterstaining with DAPI. Images were acquired using confocal microscopy to quantify the spatial distribution of proliferating (EdU+) cells within the organoid structure.

### Tissue preparation for multiome sequencing

Multimodal sequencing was performed on two distinct experimental cohorts: a time-course series of neuroids collected at days 0, 3, 4, and 7 post-differentiation, and a microenvironment reconstruction series harvested on day 7 comprising Control (CTL), Microglia (MG), Fibroblast (Fibro), and co-culture (MG_Fibro) conditions. To achieve adequate cell yields, multiple organoids from each condition were pooled and dissociated into single-cell suspensions using Accutase with EDTA, followed by 10 minutes of gentle trituration. After filtration and viability assessment via Trypan Blue, cells were cryopreserved at −80°C in CryoStor CS10 (STEMCELL Technologies, Cat. No. 07952). Libraries for simultaneous chromatin accessibility and gene expression profiling were constructed using the Chromium Single Cell Multiome ATAC + Gene Expression kit (10x Genomics). Upon rapid thawing (37°C water bath, 1–2 min), nuclei were isolated according to the 10x Genomics low-input demonstrated protocol (CG000169 Rev D), utilizing a 3-minute lysis step in a buffer containing 0.1% Tween-20 and digitonin. Subsequent single-nucleus partitioning and library preparation adhered strictly to the manufacturer’s guidelines.

Time course series multiome was sequenced using NovaSeq SP-100 cycles (whole flow cell for Gene expression library and another flow cell for the ATAC library, sequenced with NGI Sweden). Microenvironment reconstruction series multiome was sequenced using MGI DNBSEQ-G400 on separate flow cells for Gene expression and ATAC libraries (sequenced with Xpress Genomics AB).

### Processing of Neuroid scRNA-seq data: time-course analysis

Single-cell transcriptomic data from neuroids harvested at days 0, 3, 4, and 7 were processed using cellranger-arc-2.0.2 and then using Scanpy^32^ (v1.12). Quality control filters excluded cells with fewer than 100 detected genes or mitochondrial content exceeding 5%, as well as genes expressed in fewer than three cells. Data were log-normalised, and highly variable genes were selected based on expression mean (0.0125–3.0) and dispersion (>0.5). To mitigate technical variation, we regressed out confounding factors including mitochondrial and ribosomal gene content and total gene counts. Dimensionality reduction was performed via principal component analysis (PCA), retaining the top 30 components. Batch effects across time points were corrected using Harmony^33^. Clustering was conducted on a k-nearest neighbors (KNN) graph using the Leiden algorithm (resolution = 0.6), and results were visualized using Uniform Manifold Approximation and Projection (UMAP). Transcriptional programs were inferred using amortized Latent Dirichlet Allocation (LDA) via the scvi-tools framework^34^, identifying seven distinct topics. Differentially expressed genes (DEGs) were determined for each topic and cluster, with functional enrichment analysed against GO_Biological_Process_2021, CellMarker_2024, and PanglaoDB_Augmented_2021. Additionally, a scVI variational autoencoder^35^ was trained to model latent cell-state variability; high-confidence markers were defined by a mean log fold change >0 and a Bayes factor ≥2.5.

### Processing of organoid scRNA-seq data: microenvironment reconstruction

#### Data integration and preprocessing

Single-cell RNA-seq datasets from Control (CTL), Microglia (MG), Fibroblast (Fibro), and Microglia-Fibroblast co-culture (MG_Fibro) conditions were processed using cellranger-arc-2.0.2 and integrated using Scanpy (v1.12). Quality control removed cells with >20% mitochondrial content or <200 detected genes, yielding 8,071 high-quality cells. Data were normalised to 10,000 counts per cell, log-transformed, and regressed against mitochondrial percentage and gene counts. Highly variable genes were identified, and PCA was performed on scaled data (clipped to a maximum value of 10). Batch effects arising from experimental conditions were corrected using the Batch Balanced K-Nearest Neighbors (bbknn) algorithm. A UMAP embedding was generated from the bbknn-corrected neighbor graph, and unsupervised clustering was performed using the Leiden algorithm (resolution = 0.7). Cell cycle phases (S and G2/M) were scored to assess proliferation biases. Cluster markers were identified via logistic regression, while condition-specific transcriptional changes were resolved through pairwise Wilcoxon rank-sum tests comparing MG, Fibro, and MG_Fibro groups against the CTL reference within each cluster.

#### Trajectory inference and lineage reconstruction

To map the developmental landscape, trajectory inference was performed using Slingshot^36^ on the integrated data.

To reconstruct differentiation trajectories within neural lineage populations, trajectory inference was performed using Slingshot (v2.0) in R version 4.1.1. Prior to trajectory inference, single-cell RNA-seq data were processed using Scanpy in Python as described above. For trajectory inference, cluster identities derived from the Leiden clustering were used as input to Slingshot together with the PCA embedding, which preserves the global transcriptional manifold required for principal curve fitting. The cluster corresponding to quiescent neural stem cells (qNSCs) was specified as the starting cluster, allowing Slingshot to reconstruct developmental trajectories initialised from this progenitor state. Slingshot fits simultaneous principal curves through the cluster-based minimum spanning tree, generating lineage trajectories and assigning each cell a pseudotime value along each lineage as well as lineage probability weights, representing the likelihood of a cell belonging to a given lineage branch. Pseudotime values and lineage weights were exported from R and integrated back into the Scanpy AnnData object to enable downstream visualization and analysis. For each cell, a main pseudotime value was defined as the pseudotime corresponding to the lineage with the highest lineage weight.

To identify genes whose expression varied along differentiation trajectories, Spearman rank correlation was computed between gene expression and pseudotime across cells with finite pseudotime values. P-values were corrected for multiple testing using the Benjamini-Hochberg false discovery rate (FDR) procedure. Genes with FDR < 0.05 were considered significantly associated with pseudotime. For visualization of transcriptional programs along trajectories, cells were ordered by pseudotime and aggregated into quantile-based pseudotime bins (n = 25–30 bins). Mean gene expression per bin was calculated and z-score normalised per gene to highlight dynamic expression patterns. Heatmaps of the top pseudotime-associated genes were generated with hierarchical clustering of genes while preserving the pseudotime order of bins. To resolve lineage-specific transcriptional programs, pseudotime analyses were also performed separately for each lineage, restricting the analysis to cells with high lineage assignment probability (lineage weight ≥ 0.6).

To determine whether specific differentiation trajectories were preferentially utilized under different experimental conditions, lineage usage was quantified using Slingshot lineage weights. Cells were considered confidently assigned to a lineage if their lineage weight exceeded 0.6, reflecting strong membership in that trajectory. For each experimental condition (CTL, Fibro, MG, MG-Fibro), the fraction of cells assigned to each lineage was calculated relative to the total number of cells in that condition. To account for differences in total cell numbers between conditions, lineage enrichment was reported as the fraction of cells within each condition contributing to a given lineage. To visualize lineage redistribution between conditions, lineage fractions were compared to the control condition by computing: Δ lineage fraction = *f*(condition) - *f*(CTL). These differences were visualized using a diverging heatmap, highlighting trajectories enriched or depleted relative to control.

#### Topic Modeling

To dissect the gene programs underlying lineage shifts, we applied probabilistic LDA topic modeling using scvi-tools. This unsupervised approach decomposed the high-dimensional expression matrix into seven distinct transcriptional topics, each representing a coherent cellular state or functional program.

#### Ligand-receptor interaction analysis

To identify microenvironmental drivers of lineage specification, cell-cell communication was analysed using NicheNet^37^ . To identify microglia-derived signals acting on neural stem and progenitor populations, NicheNet was applied to the log-normalised single-cell RNA-seq expression matrix using the mouse ligand - target prior and ligand-receptor network. Microglia were defined as the sender population, and qNSCs, dividing NSCs, dividing cells, astrocyte precursor/radial glial-like cells, perivascular astrocytes, NPCs, astrocytes, mature astrocytes, immature neurons, neurons, OPCs, and oligodendrocytes were analysed as receiver populations. Ligand activity was evaluated for MG vs CTL and MG-Fibro vs CTL by ranking candidate ligands according to their predicted ability to explain receiver transcriptional changes, quantified using corrected Area Under the Precision-Recall curve (AUPR) scores. For prioritized ligands, corresponding ligand-receptor interactions and weighted ligand-target links were extracted from the NicheNet prior model. To assess whether microglial signalling preferentially promoted astrocytic versus neuronal programs, astrocyte-enriched and neuron-enriched gene sets were defined from the integrated dataset and used as custom receiver programs for qNSCs, dividing NSCs, NPCs, and astrocyte precursor/radial glial-like cells. Ligand activity was then calculated separately for the astrocytic and neuronal programs, and an astrocyte-minus-neuron activity score was derived for each receiver population and condition. Positive values indicate signalling biased toward astrocytic programs, whereas negative values indicate signalling biased toward neuronal programs.

## Results

### Generation and Molecular Characterisation of Adult Spinal Cord Neural Stem Cell–Derived Neuroids

A physical insult to the mammalian spinal cord activates endogenous neural stem cells (eNSCs) derived from spinal cord ependymal cells^4,6,38^. Harvested spinal cord eNSCs can be expanded in neurosphere culture^39^, where single cells clonally expand to form spheres. Although neurospheres can initially arise from progenitors with limited self-renewal capacity, sustained neurosphere formation upon serial passaging is known to be restricted to ependymal cell-derived neural stem cells (e.g., FoxJ1-CreER; tdTomato lineage traced, Fig. S1A)^4,6^. We adapted this approach to generate a spinal cord eNSC-derived spheroid/organoid, which we term a neuroid, to study neural stem cell responses, differentiation, and multicellular interactions.

In brief, spinal cord eNSCs were activated by dorsal funiculus incision in adult mice, and injured spinal cords were harvested 2 days later (Fig. 1A). Spinal cords were dissociated and cultured in neurosphere proliferation medium containing Fgf2 and Egf for 10 days to expand NSCs derived from spinal cord ependymal cells. Neurospheres were then dissociated and seeded into 96-well plates (5,000 cells per well) (Fig. 1A–B), leading to the formation of cell islands that rapidly self-organised. Individual neuroids were maintained in proliferation medium for 7 days, during which they exhibited linear growth. The medium was then exchanged to differentiation medium (Fig. S1B–C). Neuroids continued to grow in differentiation medium, reaching a plateau 3 days after induction, followed by a gradual decrease in size (Fig. S1D). Within the 3D environment, NSCs spontaneously differentiated into multiple lineages, including neurons (β-III tubulin+ detected by Tuj1 antibody) and astrocytes (Gfap+) (Fig. 1C). We did not observe a pronounced oligodendrocyte differentiation under these culturing conditions.

**Figure 1.**
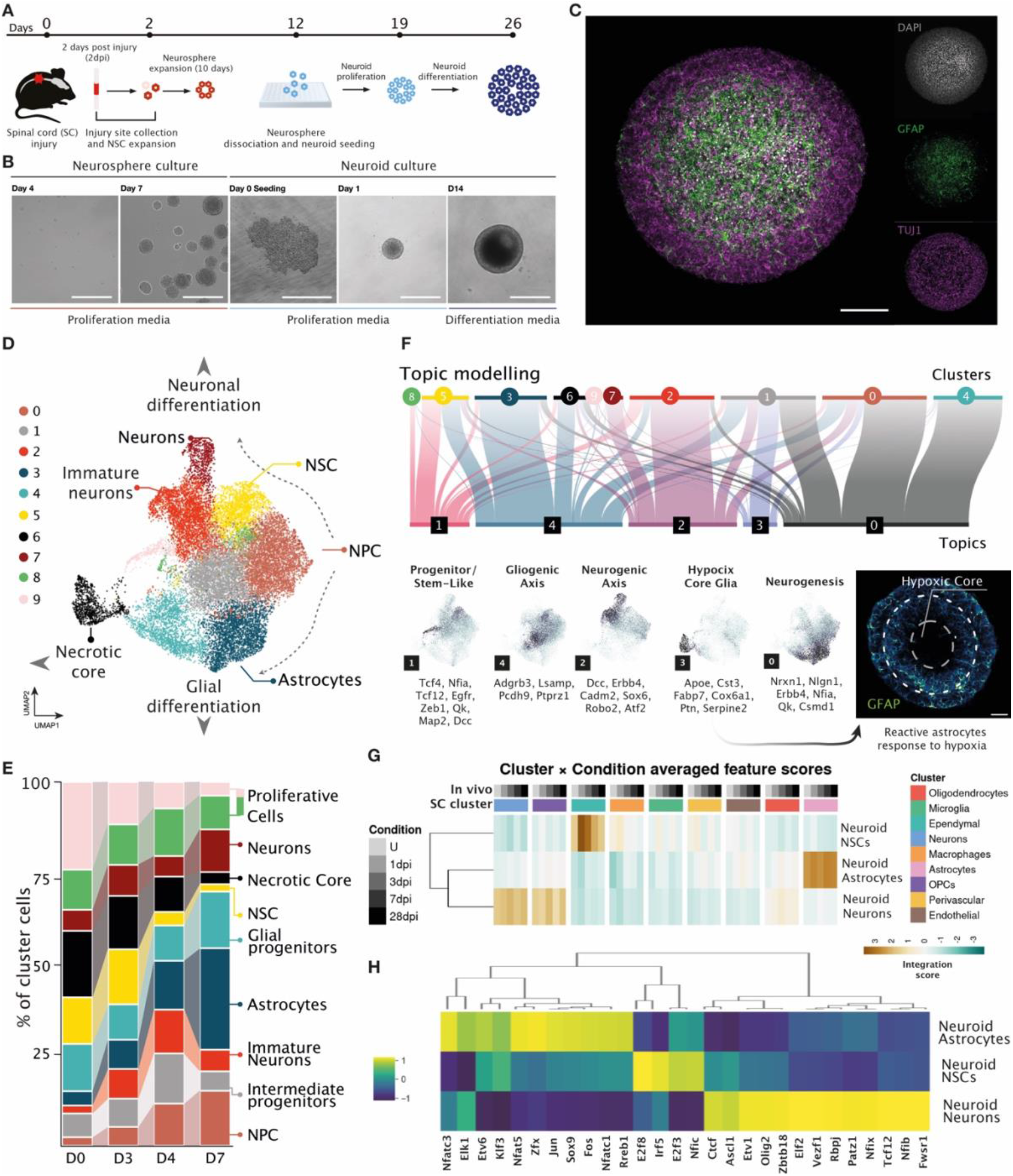
Adult Spinal Cord Neural Stem Cell-Derived Neuroid Characterisation. **(A)** Schematics depicting extraction of endogenous spinal cord neural stem cells after an injury to form neuroids. **(B**) Brightfield images of neuroid formation at different stages. Scale bar = 200 μm. **(C)** A confocal image of a differentiated neuroid at day 7 post differentiation induction stained for nuclei (DAPI in grey), astrocyte marker (GFAP in green), neuronal marker (beta-III tubulin (TUJ1 antibody) in magenta). Scale bar = 100 μm. **(D)** Uniform manifold approximation and projection (UMAP) showing identified integrated clusters from neuroids at day 0, 3, 4 and 7 after differentiation induction. **(E)** Stacked barplots separated by day of differentiation (X-axis) revealing cluster proportion (Y-axis) composition. **(F)** Top - alluvial plot connecting identified topics using topic modelling with the Leiden clusters. Bottom left - topics visualised in the UMAP space and gene lists associated with them. Bottom right - a confocal image of an organoid stained for astrocytic marker (GFAP, cyan) showing reactive astrocyte response surrounding the hypoxic neuroid core. Scale bar = 50 μm. **(G)** Heatmap of neuroid neural stem cell (NSC), astrocyte and neuron cell integration scoring based on gene expression with in vivo spinal cord cells (U – uninjured, 1, 3, 7, 28 days post injury (dpi)). **(H)** Heatmap showing uncovered enriched transcription factors based on enriched transcription factor binding sites within open neuroid astrocyte, NSC and neuron chromatin using ChromBPNet.

To further dissect neuroid composition, we collected cells at one (D1), three (D3), four (D4), and seven (D7) days after differentiation induction and performed 10x Genomics single-nuclei multiome (RNA+ATAC-seq). Dimensionality reduction, latent space transformation, and clustering analyses revealed that differentiating neuroids contain proliferating NSCs (Pola1, Kntc1, Atad2), neural progenitor cells (NPCs; Dcx), immature neurons (Dcx, Sema5a), neurons expressing mature markers (e.g., Rbfox3 [NeuN]), and astrocytes (Slc7a11, Appl2) (Fig. 1D; Fig. S1E). The NSC pool decreased from D0 to D7 (Fig. 1E) but remained detectable within differentiating neuroids. This was further supported by the ability to re-establish and passage neurospheres from dissociated differentiated neuroids, confirming the persistence of functional NSCs (Fig. S1F). Pools of astrocytes, NPCs and neurons increased over the differentiation time, while immature neuron numbers peaked at D4 followed by a decrease (Fig. 1E). We identified a cluster termed “necrotic core,” a known limitation of organoids arising from restricted oxygen and nutrient diffusion to the inner regions of densely packed 3D structures. Against the expectations, we observed a decrease in cells belonging to the necrotic core cluster over time (Fig. 1E). We attribute this to a technical quality control (QC) effect, as dying cells at later time points likely fail sequencing QC filters and are therefore excluded from downstream analyses.

We performed topic modelling using scvi-tools^34^ to identify distinctive cell states/themes shared across different cell types. This resulted in 5 distinctive topics (Fig 1F). Topic 0 was associated with neurogenesis and primarily encompassed NPC and astrocyte clusters. Topic 1 represented a progenitor/stem-like state, exhibited mainly by NSCs, as well as subsets of astrocytes, immature neurons, and necrotic core cells. Topic 2 defined a neurogenic axis including subsets of NSCs, NPCs, immature neurons, neurons, and necrotic core cells. Topic 3 corresponded to a hypoxia/core glia program associated with necrotic core cells, intermediate progenitor cells, and some NPCs. Topic 4 indicated a gliogenic axis, primarily involving NSCs, astrocytes, and necrotic core clusters.

We integrated the neuroid single cell multiome data with the in vivo spinal cord injury single cell multiome data^40^ and scored the similarity between the two data sets, which at the transcriptomic level indicated that neuroid NSCs showed highest similarity to in vivo ependymal cells, their cells of origin, neuroid astrocytes were the most similar to the in vivo astrocytes and neuroid neurons showed the most similarity to in vivo OPCs and neurons (Fig. 1G). At the chromatin level, neuroid NSCs showed similarity to in vivo OPCs; neuroid astrocytes to in vivo ependymal cells and astrocytes; and neuroid neurons to in vivo ependymal cells (Fig. S1G). We further applied ChromBPNet^41^, a deep-learning model of base-resolution chromatin accessibility, to identify regulatory transcription factor (TF) motifs associated with NSC maintenance and differentiation toward astrocytic or neuronal lineages (Fig. 1H). This revealed that neuroid NSCs are regulated by factors such as Irf5 and Nfic; astrocytes - by Sox9 and Nfatc3 among others; neurons - by Nfix, Etv1, Ascl1 and others.

We also tested neuroid culture in Matrigel, however, this resulted in excessive cell migration and loss of structural integrity (Fig. S1H). We therefore continued culturing neuroids in suspension.

### Reconstitution of Fibrotic Scar Architecture within Neuroids

Given that the spinal cord injury environment is highly pro-glial and anti-neurogenic - where ependymal cell-derived NSCs primarily differentiate into glial scar-forming astrocytes, rarely into oligodendrocytes, and never into neurons ^6,42^ - we aimed to establish a bottom-up neuroid system with a modularly reconstituted spinal cord injury microenvironment to investigate how environmental cues influence NSCs and their multilineage progeny.

We first reconstituted fibrosis by adding lineage-traced meningeal fibroblasts (Col1a1-CreERT2-TdTomato+) after induction of neuroid differentiation (Fig. 2A). As early as day 1 after addition, fibroblasts began populating the interior of the neuroids (Fig. 2B), predominantly localizing to the neuroid core (Fig. 2C). We additionally incorporated sorted fibroblasts reporting active collagen I expression (Col1a1-GFP+). This revealed that fibroblasts within the neuroid core actively produce collagen, indicating functional extracellular matrix deposition in situ (Fig. 2D).

**Figure 2.**
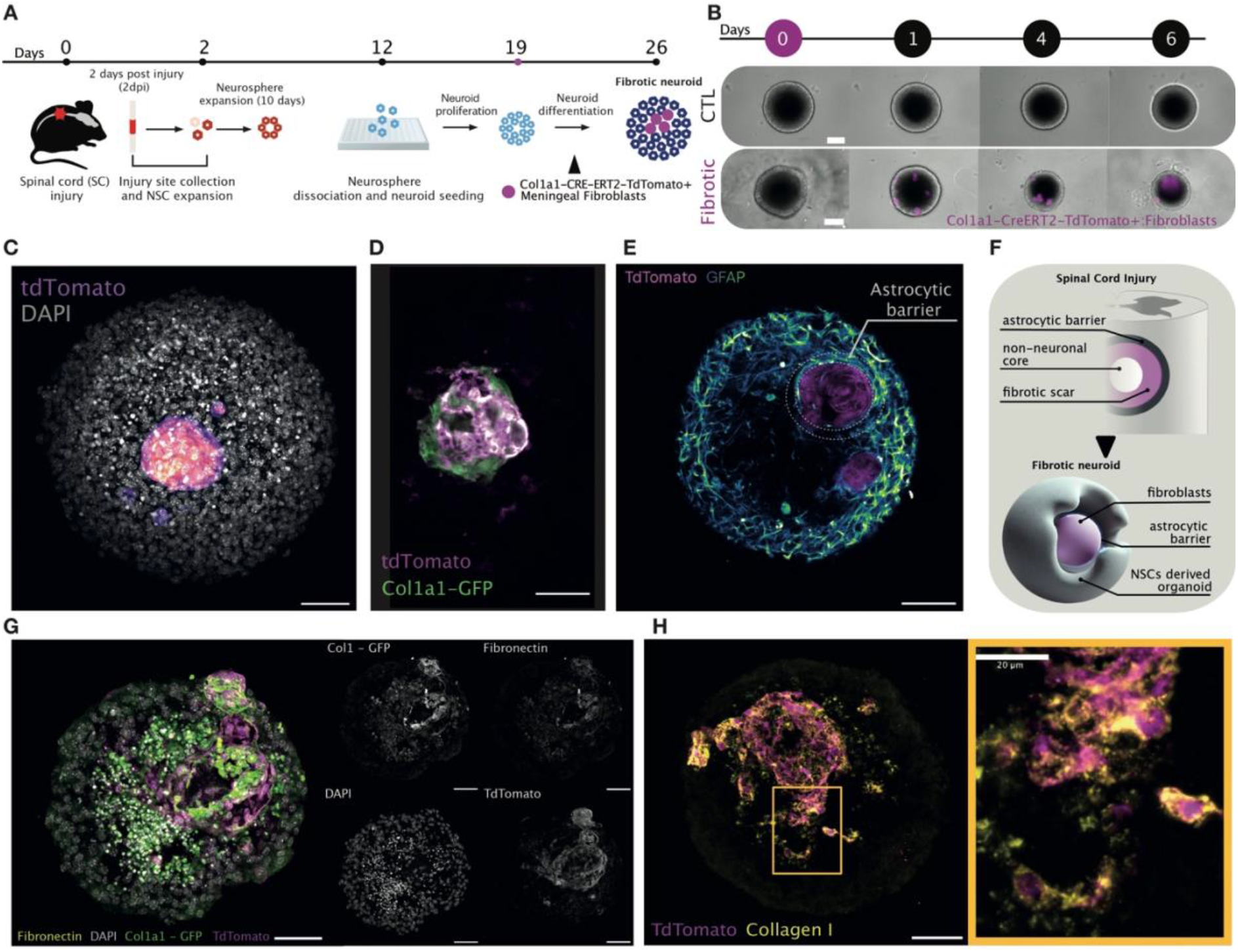
Fibrosis Reconstitution Within Neuroids. **(A)** Schematics depicting meningeal fibroblast incorporation within neuroids. **(B)** Brightfield images showing fibroblast (in magenta) infiltration into neuroids at day 0, 1, 4 and 6 post seeding. Scale bar = 200μm **(C)** A confocal image of a differentiated neuroid at day 7 post differentiation induction after fibroblast infiltration stained for nuclei (DAPI in grey) and tdTomato to visualise seeded Col1a1-CreERT2-tdTomato fibroblasts (in magenta). Scale bar = 50 μm. **(D)** A confocal image of a fibrotic neuroid core with lineage traced Col1a1-CreERT2-tdTomato fibroblasts (in magenta) and fibroblasts actively reporting collagen 1 expression (Col1a1-GFP, in green). Scale bar = 50 μm. **(E)** A confocal image of a differentiated neuroid at day 7 post differentiation induction after fibroblast infiltration showing an astrocytic border (highlighted by dashed circles, GFAP in green) around the fibrotic core (tdTomato+ fibroblasts in magenta). Scale bar = 50 μm. **(F)** Schematics depicting similarities between in vivo spinal cord injury formed scar and reconstituted fibrotic neuroid scar architectures. **(G)** Left - a confocal image of fibrotic neuroid stained for extracellular matrix protein fibronectin (in yellow), fibroblasts reporting collagen 1 expression (Col1a1-GFP in green, lineage traced meningeal fibroblasts (Col1a1-CreERT2-tdTomato, in magenta), nuclei (DAPI in grey). Right - separated channels for each stained protein of the image on the left, signal visualised in grey. Scale bar = 50 μm. **(H)** Left - a confocal image of fibrotic neuroid stained for extracellular matrix protein collagen 1 (in yellow and lineage traced meningeal fibroblasts (Col1a1-CreERT2-tdTomato, in magenta). Scale bar = 50 μm. Right - a zoomed in view of a fibrotic neuroid area boxed in yellow.

We tested different numbers of fibroblasts and varied the timing of addition, including during the neuroid proliferation stage (Fig. S2A–B). The size of the fibrotic core could be modulated by the initial number of fibroblasts added. For example, addition of 10,000 fibroblasts resulted in large lesion-like structures (Fig. S2C). We concluded that adding 500-1000 TdTomato+ sorted fibroblasts to differentiating neuroids was optimal for preserving structural integrity. This condition still resulted in an overall reduction in neuroid size compared to control neuroids (Fig. S2D), possibly due to increased compaction and tissue stiffening.

Fibroblasts populating the neuroid core proliferated (Fig. S2E) and induced reactivity in adjacent astrocytes, mimicking the in vivo scar architecture in which a fibrotic core is surrounded by an astrocytic barrier (Fig. 2E–F). We further evaluated ECM production by fibrotic neuroid core fibroblasts using fluorescent immunohistochemistry for fibronectin (Fig. 2G) and collagen I (Fig. 2H) proteins. Added fibroblasts colocalized with ECM protein staining, confirming their functional contribution to in vitro fibrotic core formation. We did not observe any spontaneous ECM protein deposition in control neuroids (Fig. S2F).

To assess alternative fibroblast sources, we tested primary skeletal muscle fibroblasts and an embryonic mouse fibroblast cell line (Fig. S2G–H). However, these cells behaved differently and largely overgrew the neuroid structures. We therefore continued using primary meningeal fibroblasts as the most physiologically relevant source.

### Reconstitution of the Innate Immune Microenvironment within Neuroids

Spinal cord injury leads to rapid activation of resident immune cells, microglia, establishing an inflammatory microenvironment^43^. Such environments have been linked to limited neurogenesis and reinforced gliogenic programs in the brain^15,16^, however, how inflammatory signalling regulates spinal cord NSCs after injury remains incompletely understood. To model this critical component of the injury niche, we integrated microglia into neuroids to reconstitute the innate immune compartment and interrogate how inflammatory cues instruct NSC fate decisions.

To this end, we isolated primary adult microglia from brain tissue and initially co-cultured them with astrocytes using an established isolation protocol^9^. Microglia were subsequently detached from the astrocyte layer, re-cultured, and then added to differentiating neuroids in suspension (Fig. 3A; Fig. S3A). Cultured adult microglia expressed Cd45 and Ki67, confirming their identity and proliferative capacity in vitro (Fig. 3B). We further characterized cultured microglia by flow cytometry to confirm the purity of cells added to neuroids (Fig. S3B’, B’’). Following addition to the suspension culture, microglia rapidly localized to neuroids within hours and populated their interior (Fig. 3C). In contrast to fibroblasts, microglia did not concentrate within the neuroid core but instead exhibited a more dispersed and heterogeneous distribution (Fig. 3D). No microglial markers were detected in control neuroids, indicating the absence of spontaneous microglial differentiation, consistent with their distinct developmental origin^44^ (Fig. S3C).

**Figure 3.**
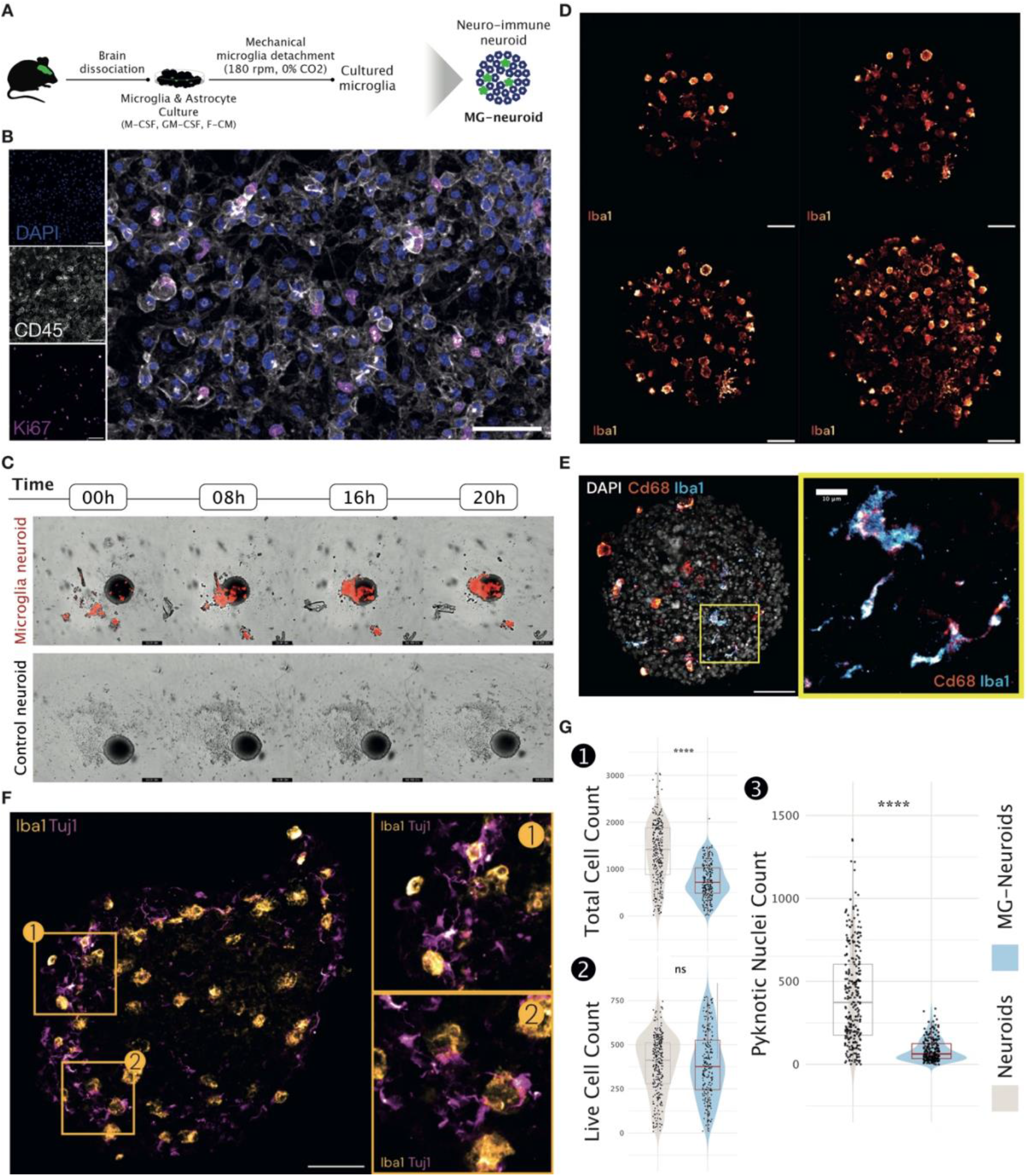
Tissue Resident Microglial Cell Reconstitution Within Neuroids. **(A)** Schematics depicting acquisition and culturing of primary adult microglial cells. **(B)** Confocal image of passaged and cultured microglial cells in monolayer stained for hematopoietic Cd45 cell marker (in grey), nuclei (DAPI in blue) and cell division marker Ki67 (in magenta). Scale bar = 50 μm. **(C)** Brightfield images showing microglia (stained in red) infiltration into neuroids at day 0, 8, 16 and 20 hours post seeding. **(D)** Confocal images of neuroid sections infiltrated with microglia (Iba1 in yellow-red). Scale bar = 50 μm. **(E)** Left - a confocal image of neuroid section with infiltrated microglia detected by phagocytic (Cd68, in red) and homeostatic (Iba1, in blue) microglial markers. Scale bar = 50μm. Right - a zoomed in view of a microglia populated neuroid area boxed in yellow. **(F)** Left - a confocal image of a neuroid section detecting microglia (Iba1, in yellow) and neurons (beta-III tubulin (Tuj1 antibody) in magenta). Scale bar = 50 μm. Right - zoomed in view of a microglia populated neuroid two areas boxed in yellow highlighting microglia-neuron interactions. **(G)** 1 – Violin plot showing total cell count in control neuroids and microglia populated neuroids (MG-neuroids). 2 - Violin plot showing live cell count in control neuroids and MG-neuroids. 3 - Violin plot showing pyknotic nuclei count in control neuroids and MG-neuroids. Statistical significance was assessed using a two-sided Wilcoxon rank-sum test. **** *p < 0*.*0001*; ns, not significant.

Integrated microglia expressed both phagocytic (Cd68) and homeostatic (Iba1) markers and displayed either activated/phagocytic or ramified morphologies, suggesting the coexistence of activated/pro-inflammatory and homeostatic states within neuroids (Fig. 3E). We observed direct interactions between microglia and neurons, indicating that integrated microglia are functionally active and may contribute to synaptic pruning, remodelling, and neuronal homeostasis maintenance (Fig. 3F).

In addition to Cd68 staining, we further validated the phagocytic capacity of microglia within neuroids. Addition of microglia resulted in a significantly lower neuroid cell count compared to control neuroids, however, total live cell numbers were not affected, suggesting that microglia facilitate removal of dead cells (Fig. 3G). Neuroids containing microglia exhibited significantly fewer pyknotic nuclei compared to controls, and engulfed pyknotic nuclei were observed within Iba1+ cells (Fig. 3G; Fig. S3D). Consistent with this, apoptosis was reduced in microglia-containing neuroids, as assessed by Cleaved Caspase-3 (CC3) staining (Fig. S3E).

Together, these findings demonstrate that microglia integrate functionally within neuroids, adopt heterogeneous activation states, and actively participate in cellular remodelling of the 3D neural environment.

### Reconstruction of a Composite Fibrotic–Inflammatory Injury Niche

In vivo, spinal cord injury establishes a complex lesion environment in which fibrotic scar formation and microglial activation coexist and dynamically interact. Following injury, activated microglia produce a range of cytokines, including TGFβ, a key mediator of fibroblast recruitment, activation, and differentiation into myofibroblasts, thereby contributing to fibrosis^22,45^. However, dissecting the individual and combinatorial contributions of these niche components to neural stem cell behaviour and lineage decisions remains experimentally challenging in vivo. We therefore sought to reconstruct a composite fibrotic-inflammatory microenvironment within neuroids by incorporating both meningeal fibroblasts and microglia and to evaluate how this more complex in vitro injury niche influences NSC differentiation trajectories.

To achieve this, we first added microglia at the onset of neuroid differentiation and allowed them to populate the neuroids. Two days later, meningeal fibroblasts were incorporated using the same approach as described above (Fig. 4A). This recapitulated the previously observed seeding patterns: microglia dispersed throughout the neuroid, whereas fibroblasts accumulated within the neuroid core (Fig. 4B).

**Figure 4.**
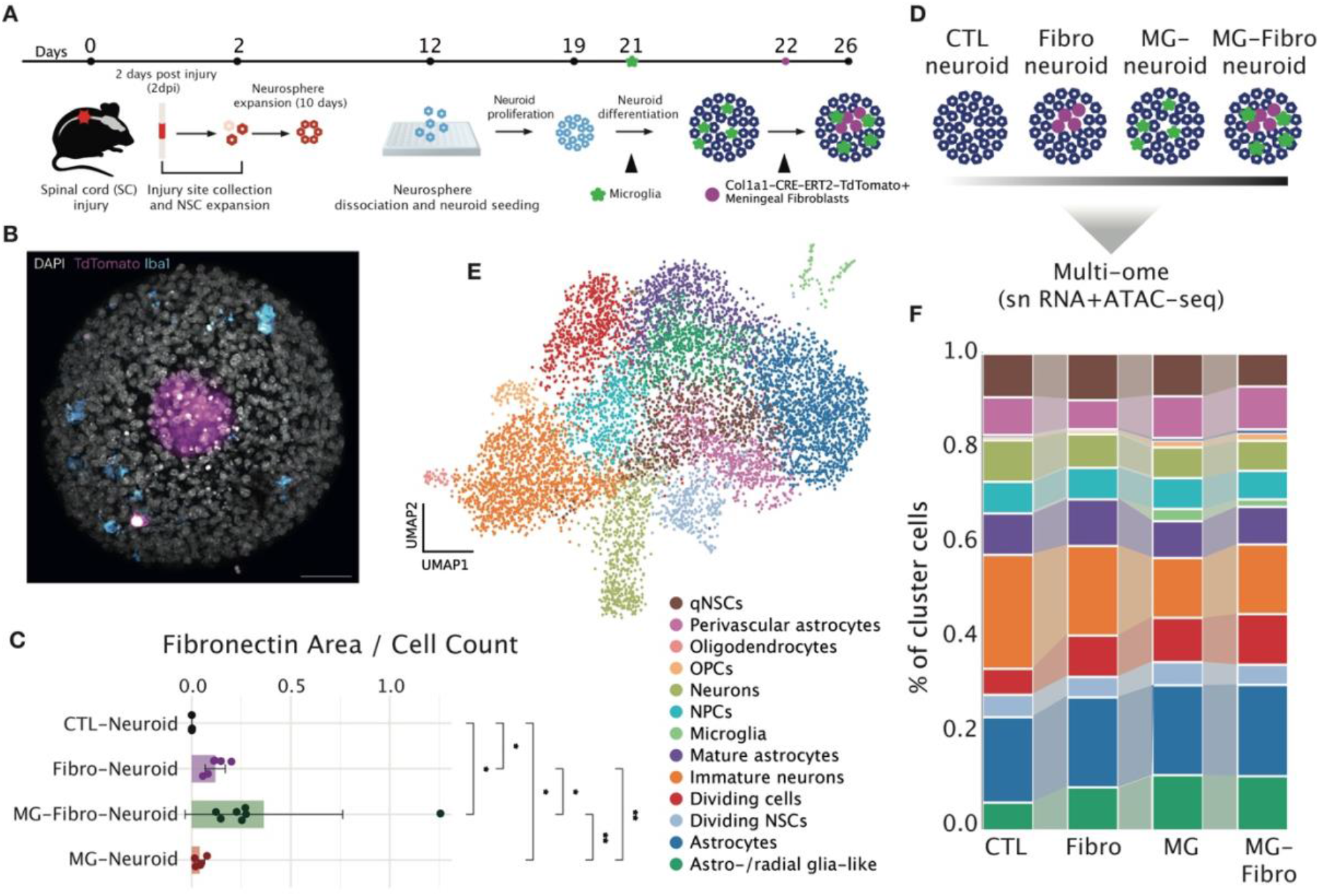
Composite Fibrotic-Inflammatory Injury Niche Reconstitution Within Neuroids. **(A)** Schematics depicting adult microglia and meningeal fibroblast incorporation within neuroids. **(B)** A confocal image of a differentiated neuroid at day 7 post differentiation induction after microglia and fibroblast infiltration stained for nuclei (DAPI in grey), Iba1 (in cyan) and tdTomato (to visualise seeded Col1a1-CreERT2-tdTomato fibroblasts, in magenta). Scale bar = 50 μm. **(C)** A bar plot showing quantified fibronectin area (measured by staining against fibronectin) per cell count (X-axis) in different neuroid conditions (Y-axis: CTL - control, Fibro neuroid - neuroids with fibroblasts, MG-neuroid - neuroids with microglia, MG-Fibro neuroid - neuroids with both incorporated microglia and fibroblasts. Error bars represent mean ± standard deviation, with individual data points overlaid (n = 2-6 neuroids per condition). Statistical significance was evaluated using a pairwise Wilcoxon rank-sum tests with Bonferroni correction. Significance levels are indicated as: *\*p < 0*.*05, **p < 0*.*005*. **(E)** Uniform manifold approximation and projection (UMAP) showing identified integrated clusters from control, Fibro-, MG- and MG-Fibro neuroids at 7 days after differentiation induction. **(F)** Stacked barplots separated by neuroid condition (X-axis) revealing cluster proportion (Y-axis) composition. Same cluster annotation colours as in E.

Interestingly, addition of microglia alone led to detectable fibronectin deposition within neuroids, albeit to a lesser extent than in fibrotic neuroids (Fig. 4C). In contrast, combined addition of microglia and fibroblasts resulted in a significant increase in fibronectin deposition compared to fibrotic neuroids alone (Fig. 4C; Fig. S4A), suggesting cooperative interactions between microglia and fibroblasts that enhance extracellular matrix accumulation and generate a more heterogeneous scar-like architecture. We further observed that enhanced fibrosis was accompanied by innervation of the scar-like regions by neuroid-derived neurons (Fig. S4B).

We next applied 10x Genomics single-nuclei multiome profiling to compare control (CTL), fibrotic (Fibro), microglia-containing (MG), and combined fibrotic-microglial (MG-Fibro) neuroids at D7 after differentiation induction (Fig. 4D). Following dimensionality reduction and clustering analyses, a distinct fibroblast cluster was not detected (Fig. 4E). This likely reflects the rigid extracellular matrix environment, from which fibroblasts may not have efficiently dissociated into single-cell suspensions for downstream processing. We identified a microglial cluster exhibiting a phagocytic signature (Cd68+, Ptprc+ [Cd45], Itgam+ [Cd11b]) that was present exclusively in MG and MG-Fibro neuroids, as expected (Fig. 4E– F; Fig. S4C–E).

Notably, we detected pronounced oligodendrocyte progenitor cell (OPC) and oligodendrocyte clusters in this dataset (Fig. 4E; Fig. S4C–E), which were not evident in the previously analysed neuroids lacking a reconstructed microenvironment (Fig. 1D). Although a small number of CTL-derived cells mapped to this lineage, oligodendrocyte-lineage cells were predominantly observed in Fibro, MG, and particularly MG-Fibro neuroids (Fig. S4C). These findings suggest that the reconstituted injury-like environment permits or enhances oligodendrocyte lineage commitment.

Integration of all samples revealed 13 clusters, including quiescent NSCs (qNSCs), dividing NSCs, cycling progenitors, neural lineage cells, and astrocyte-lineage populations (Fig. 4E; Fig. S4E). Neuroids containing fibroblasts and/or microglia exhibited increased proportions of dividing cells and astro-/radial glia-like populations (Fig. 4F). In contrast, CTL neuroids were enriched for immature neuron and mature neuron clusters.

Together, these data indicate that neurogenesis is putatively more permissive in CTL neuroids, whereas incorporation of injury-associated niche components shifts lineage composition toward glial and progenitor states, thereby reshaping NSC differentiation trajectories (Fig. 4F).

### Injury-Like Microenvironment Biases Neural Stem Cell Lineage Trajectories Toward Gliogenesis

The observed changes in cell cluster proportions between control neuroids and neuroids with reconstituted injury niche components prompted us to examine whether reconstructed microenvironments influence neural stem cell differentiation trajectories at a systems level.

To this end, we first reconstructed cellular trajectories in neural organoids using lineage inference and pseudotime analysis in Slingshot^36^. Lineage relationships and pseudotime were inferred based on PCA embeddings, using qNSCs as the starting cluster (Fig. S5A–B), and trajectories were subsequently visualized in UMAP space (Fig. 5A). This resulted in six inferred differentiation lineages: Lineage 1 - transitioning through dividing cells toward OPCs/oligodendrocytes and immature neurons; Lineage 2 - progressing toward immature neurons via NPC intermediates; Lineage 3 - NPCs differentiating toward mature neurons; Lineage 4 - passing through astro-/radial glia-like states toward astrocytes and mature astrocytes; Lineage 5 - transitioning toward dividing NSCs; and Lineage 6 - differentiating toward astrocytes.

**Figure 5.**
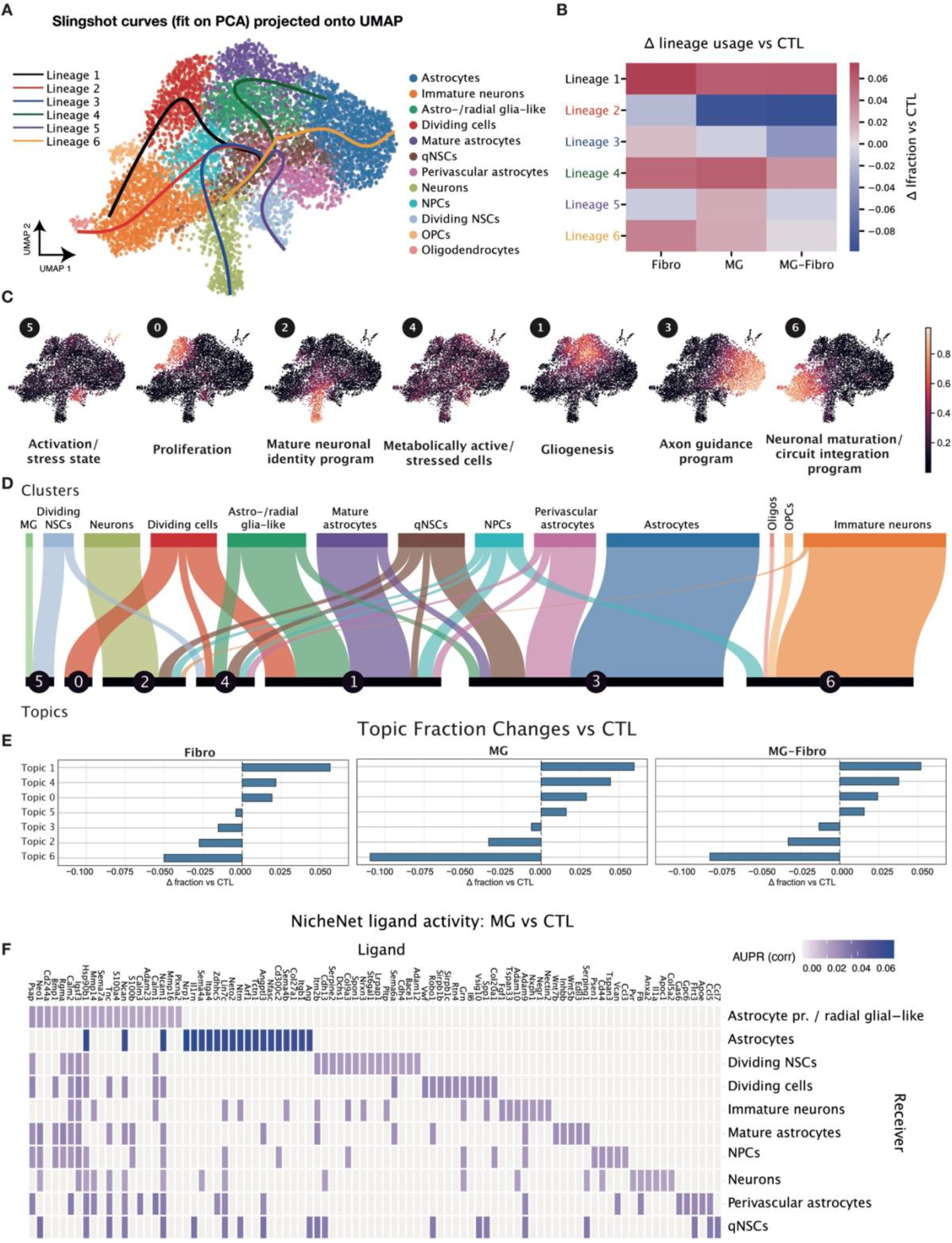
Recapitulation of in vivo Neural Stem Cell Lineage Biases Within Injury-Like Reconstituted Microenvironment. **(A)** Cell lineage trajectories uncovered by Slingshot algorithm inferred based on PCA embeddings and visualised on uniform manifold approximation and projection (UMAP) space. **(B)** Heatmap showing the change in the fraction of cells assigned to each Slingshot-inferred lineage relative to control (CTL). Values represent the difference in lineage usage (Δ fraction vs CTL) for fibrotic (Fibro), microglia seeded (MG) and microglia and fibroblast (MG-fibro) containing neuroid conditions. Positive values (red) indicate increased representation of a lineage compared with CTL, whereas negative values (blue) indicate reduced representation. **(C)** Identified topics using topic modelling visualised in the UMAP space. **(D)** Alluvial plot connecting identified topics using topic modelling with annotated Leiden clusters. **(E)** Bar plots showing topic fraction changes in Fibro, MG and MG-fibro neuroid data compared to the control neuroids. **(F)** Heatmap showing NicheNet-predicted ligand activity for microglia-derived ligands across neuroid receiver populations in the MG-neuroid vs CTL-neuroid comparison. Ligands are shown on the X-axis and receiver cell types on the Y-axis. Color intensity represents the corrected ligand activity score (AUPR), indicating the predicted strength of ligand-driven transcriptional responses in each receiver population.

We next examined gene expression dynamics along lineage pseudotime (Fig. S5C). For instance, along the qNSC-to-neuron trajectory (lineage 3), early pseudotime was enriched for stem and progenitor-associated regulators including Egfr, Notch2, Msi2, and Wnt effectors (Tcf7l1/Tcf7l2), consistent with maintenance of neural stem cell identity. As cells progressed, neuronal commitment markers such as Ntrk2, Dcc, and adhesion molecules including Ncam2, followed later by Mdga1, became upregulated, reflecting transition toward a post-mitotic neuronal state. Late pseudotime was characterized by strong induction of synaptic and excitability-related genes, including glutamate receptor subunits (Gria2, Grik2), mature neuronal markers (Rbfox3), and regulators of synaptic protein expression such as Nova1, indicative of terminal neuronal differentiation (Fig. S5C). Together, these transcriptional dynamics recapitulate canonical features of neurodevelopment, supporting the biological fidelity of both the neuroid model and the reconstructed lineage trajectories.

To assess neuroid condition-specific lineage bias, we quantified the proportion of cells assigned to each Slingshot-derived trajectory using lineage probability weights and compared lineage representation across experimental groups. This analysis revealed differential enrichment of specific trajectories between conditions (Fig. 5B). Specifically, lineage 2 (immature neuron lineage) was enriched in CTL neuroids compared to Fibro, MG and MG-Fibro neuroids. Lineage 3 (mature neuron lineage) usage was also reduced in MG-Fibro neuroids relative to CTL neuroids. These findings support our initial observations that reconstructed microenvironments are less favourable for neuronal differentiation.

Conversely, lineage 1 (transition through proliferative intermediates toward OPC/oligodendrocyte and immature neuronal states) was enriched in all microenvironment-reconstituted neuroid groups, particularly in Fibro-neuroids, compared to the control. This suggests that the reconstructed microenvironment contributes to increased cellular proliferation, mimicking injury-induced activation of cell cycling, including neural stem cell expansion. It also supports that microenvironment supports oligodendroglia lineage via the expansion of proliferating progenitors. In addition, astrocytic lineages (lineages 4 and 6) were enriched in microenvironment-reconstituted organoids compared to the control, further emphasizing that the injury-like microenvironment is more permissive for astrogliosis rather than neurogenesis. This closely mirrors the in vivo spinal cord injury scenario and supports the idea that while ependymal cell-derived NSCs possess neurogenic potential, injury-associated microenvironmental cues bias their differentiation toward glial fates.

To further examine cell state programs associated with these lineage shifts, we applied topic modelling to the integrated dataset and identified seven transcriptional topics capturing distinct cellular states and programs (Fig. 5C–D, Fig. S6A). Topic 0 was associated with proliferation and was primarily linked to the dividing cell cluster. Topic 1 represented a gliogenesis-associated program, connected to mature astrocytes, astro-/radial glia-like cells, and subsets of perivascular astrocytes, dividing cells, qNSCs and NPCs. Topic 2 corresponded to a mature neuronal identity program and was mainly associated with neuronal clusters, as well as subsets of qNSCs, NPCs and immature neurons. Topic 3 reflected an axon guidance program and was largely associated with astrocytes and perivascular astrocytes, suggesting a potential contribution of astrocytes to axonal guidance or structural organisation within neuroids. Topic 4 captured general metabolic and stress-associated features and was present across multiple clusters. Topic 5 corresponded to an activation or stress-associated state, particularly associated with microglia and dividing NSCs. Finally, Topic 6 represented neuronal maturation and circuit integration programs, enriched primarily in immature neurons, OPCs, oligodendrocytes and subsets of NPCs.

We next compared topic fractions across conditions (Fibro, MG and MG-Fibro) relative to CTL neuroids to determine how reconstructed microenvironments alter transcriptional programs (Fig. 5E). All microenvironment-reconstituted samples showed decreased fractions of Topics 2, 3 and 6, particularly in MG-Fibro neuroids. Thus, using an orthogonal analysis similarly indicated that reconstructed injury-like environments are less permissive for neurogenesis and neuronal maturation. Conversely, gliogenesis and proliferation-associated topics were enriched in microenvironment-reconstituted samples, emphasizing that injury-associated microenvironmental cues shift progenitor populations toward proliferative and glial differentiation programs.

To understand which microenvironmental factors might contribute to this shift toward gliogenic versus neurogenic outcomes, we performed NicheNet analysis to rank microglia-expressed ligands predicted to act on neuroid receiver cell populations based on downstream transcriptional responses (Fig. 5F, Fig. S6 B-D). This approach prioritises ligands whose activity best explains gene expression changes observed in receiver cells. Because fibroblasts were not captured in the recovered single-cell dataset, we were unable to directly evaluate fibroblast-derived ligands. However, as fibroblast-microglia interactions may alter microglial signalling states, we analysed MG-Fibro samples to determine whether microglia express distinct ligand repertoires when co-existing with fibroblasts in the reconstructed microenvironment.

In MG neuroids, predicted microglial signalling toward astrocytes was among the strongest, with Ang, Sema4a/4b, Il1rn, and Itgb1 as highly ranked candidates, alongside ECM-associated ligands Tnc, Mmp16, and Ncan predicted to act on astrocyte precursor/radial glial-like cells (Fig. 5F). Il6 was predicted to signal to dividing cells - IL-6/JAK-STAT3 is a well-established pathway promoting astrogliogenesis from neural progenitors and suppressing neuronal differentiation^46^. Predicted signalling toward qNSCs included Tnc, Ncan, Spp1, Angptl3, Il1rn, Ccl5, and Ccl7 among ranked candidates. Osteopontin/SPP1 is a known microglial-derived signal upregulated following CNS injury and linked to glial activation and reactive responses^47,48^. Tnc and Ncan are ECM glycoproteins upregulated in the spinal cord glial scar niche^49,50^ and in an indirect way might further promote gliosis due to the matrix surrounding NSCs stiffening, which is known to promote NSC differentiation into glial cells rather than neurons^51^. Moreover, Tnc was identified to reside in the stem cell niche in the subependymal zone contributing to neural stem cell activation and cell cycle progression^52^.

In MG-Fibro neuroids, predicted signalling was substantially amplified across multiple receiver populations (Fig. S6B). For astrocytes, Tgfb1, Fgf1, Sema3a, and Mmp16 were among the most highly ranked ligands. For astrocyte precursor/radial glial-like cells, predicted ligands included Tnc, Mmp16, Wnt5b, Bmp1, and Angptl3 - reflecting a broader ECM remodelling and pro-gliogenic signalling landscape compared to MG alone. For dividing NSCs, Spp1 was identified again, consistent with microglial injury-associated signalling toward glial cell states, alongside other ECM and adhesion-associated candidates. In MG-Fibro neuroids, the predicted signalling landscape toward qNSCs was further enriched, with ECM-associated candidates including Tnc and Angptl3 retained from MG-only conditions alongside an expanded chemokine repertoire - notably the addition of Ccl3 to the Ccl5 and Ccl7 predicted in MG neuroids. Ccl3, Ccl5, and Ccl7 are among the most rapidly induced signals in the injured spinal cord in vivo, with Ccl3 and Ccl7 peaking in the acute phase and Ccl5 accumulating during the subacute inflammatory phase ^53,54^, but their direct roles in qNSC regulation remain to be established.

Predicted microglial signalling toward mature neurons, while present, was comparatively lower than toward astrocytes in both conditions. In MG neuroids, ranked ligands toward neurons included ECM and structural molecules (Tnc, Ncan, Ncam1, Mmp14) alongside inflammatory and injury-associated signals (Anxa2, Il1a, Apoc1, Serping1) - consistent with a damage-response rather than pro-neurogenic signalling profile (Fig. 5B). In MG-Fibro neuroids, the predicted ligand repertoire signalling to neurons shifted toward synaptic adhesion and axonal organisation molecules including Nrxn1, Nrxn3, Nlgn2, and Nfasc (Fig. S6 B-D), suggesting that fibroblast co-presence may enhance microglial interactions involved in synaptic maintenance and circuit organisation.

Finally, we quantified this bias by scoring whether microglial ligands were associated with pro-neuronal or pro-astrocytic transcriptional programs across NSC populations. This analysis revealed that microglial ligands were predominantly associated with astrocytic programs in neuroids with reconstructed injury environments (Fig. S6E), further supporting the notion that microenvironment-derived cues actively drive NSC differentiation toward glial rather than neuronal fates.

## Discussion

In this study we developed a modular organoid system derived from adult spinal cord neural stem cells that enables reconstruction of key components of the spinal cord injury microenvironment. Using this platform, we show that incorporation of microglia and fibrotic scar-forming fibroblasts into neuroids is feasible and functional. As a result, using this microenvironment reconstituted neural stem cell nice, allowed us to uncover that it alters neural stem cell lineage trajectories, shifting differentiation away from neuronal programs and toward proliferative and astroglial states. These observations support the concept that environmental signals within the injury niche actively instruct endogenous neural stem cell fate decisions.

Previous studies uncovered that ependymal cells comprise the neural stem cell population in the adult spinal cord and become activated following injury^4–6,55^. Although these cells retain multilineage potential, their differentiation in vivo is heavily biased towards astrocyte production and glial scar formation^4,11^. Our neuroid system recapitulates this phenomenon in a controlled environment. In the absence of reconstructed niche components, neuroids display robust neuronal differentiation. However, when exposed to inflammatory and fibrotic microenvironmental cues, neuronal lineages are reduced while proliferative and astrocytic trajectories become enriched. This shift mirrors lineage outcomes observed following spinal cord injury and suggests that injury-associated signals contribute to stem cell activation/proliferation and bias neural stem cell differentiation outcomes. Notably, the observed lineage bias in our system likely represents an underestimate of a full in vivo scenario, as we reconstitute only selected components of the injury niche. The full injury environment encompasses a considerably broader range of inflammatory cells and signals, including infiltrating macrophages, T cells, and neutrophils, that may further exacerbate gliogenic bias and suppress neurogenesis. The modular nature of the neuroid system makes it suited to address this in future studies by incorporating additional niche components and dissecting their individual and combined contributions to neural stem cell fate.

The modular design of the neuroid system also allowed reconstruction of spatial features resembling the scar architecture. Seeded fibroblasts preferentially localised to the neuroid core and deposited extracellular matrix proteins including fibronectin and collagen. In spinal cord injury, fibroblasts similarly accumulate in the lesion centre and generate a fibrotic scar surrounded by reactive astrocytes^56^. Interestingly, fibroblasts consistently migrated toward the neuroid core, which likely reflects the hypoxic or necrotic core commonly observed in organoid systems^23^. Although often considered a limitation of organoid cultures, in our system it rather served as an advantage to provide a permissive niche for fibroblast recruitment and extracellular matrix deposition, thereby facilitating reconstruction of scar-like tissue organisation.

Single-cell transcriptomic analysis further revealed that reconstructed microenvironments alter neural stem cell programs at multiple regulatory levels. Lineage trajectory inference demonstrated that neuronal differentiation pathways were enriched in control neuroids, whereas astrocytic and proliferative trajectories were preferentially used in neuroids containing microglia and fibroblasts. Topic modelling supported this observation by identifying transcriptional programs associated with gliogenesis and proliferation that were enriched in injury-like conditions, while neuronal maturation programs were reduced.

Ligand-receptor analysis suggested that microglia provide signals potentially capable of influencing neural stem and progenitor populations in this system. In MG neuroids, highly ranked candidate ligands included inflammatory mediators and ECM-associated molecules linked to astrogliosis, stem/progenitor regulation, and injury-associated niche remodelling. In MG–Fibro neuroids, the predicted microglia-derived ligand list was further expanded, with enrichment of ECM-remodelling, chemokine, TGF-β-related, and non-canonical Wnt-associated candidates, consistent with a microenvironment that favours gliogenic over neurogenic programmes. It is important to note that these interpretations should be viewed in the context of NicheNet inference, which prioritises ligands whose activity best explains receiver-cell transcriptional responses rather than establishing direct ligand function. Nevertheless, the observed shift is consistent with prior work showing that stromal and immune cell interactions shape inflammatory and fibrotic responses after CNS injury^22,45,57^. In line with this, we also observed increased ECM deposition in neuroids co-seeded with both microglia and fibroblasts compared with fibroblast-only neuroids, further supporting functional microglia-fibroblast interactions in this 3D injury-like environment.

Organoid models of spinal cord injury have recently begun to emerge, including systems where spinal cord organoids are subjected to mechanical injury to model neuronal damage and inflammatory responses^58^. While such approaches capture aspects of acute trauma, they typically rely on developmentally patterned tissues derived from pluripotent stem cells. In contrast, the neuroid system described here focuses on adult neural stem cells and reconstructs the cellular microenvironment of the injury niche rather than the mechanical insult itself. These complementary approaches provide distinct opportunities to investigate injury biology: mechanical injury models capture acute tissue damage responses, whereas the neuroid system enables systematic dissection of environmental signals that regulate endogenous neural stem cell behaviour.

The modular nature of this platform also provides opportunities for future mechanistic studies. Additional cellular components of the injury niche, including endothelial cells, infiltrating immune cells, or pericytes, could be incorporated to further increase physiological complexity and relevance, and in a modular and combinatorial manner. Because the system allows controlled manipulation of environmental signals, it will also enable systematic perturbation of candidate pathways using genetic or pharmacological approaches. Such strategies may help identify environmental cues that, for instance, suppress or promote neuronal differentiation programs, while at the same time reduce fibrosis, which might pave a way for future regenerative therapies.

Finally, although the current study focuses on mouse-derived neural stem cells, the conceptual framework of this system could be extended to human cells. Integration of human neural progenitors together with immune and stromal components may provide a platform to model human spinal cord injury responses and evaluate candidate regenerative strategies, which often do not translate from animal research findings.

## Conclusions

Together, our findings establish a bottom-up organoid system that reconstructs key aspects of the spinal cord injury microenvironment and reveals how environmental cues bias endogenous neural stem cell differentiation. By enabling controlled interrogation of niche-derived signals, this platform may help identify strategies to redirect endogenous neural stem cells toward regenerative outcomes after spinal cord injury.

## Acknowledgements

The authors acknowledge support from the National Genomics Infrastructure in Stockholm funded by Science for Life Laboratory, the Knut and Alice Wallenberg Foundation and the Swedish Research Council, and NAISS for assistance with massively parallel sequencing and access to the UPPMAX computational infrastructure.

The authors thank the Biomedicum Imaging Core (BIC) facility (Karolinska Institutet) for technical expertise and assistance with imaging.

The authors acknowledge support from Xpress Genomics AB with the help of DNA library sequencing.

The authors want to thank to the Comparative Medicine Biomedicum facility dedicated to research on small rodents with invaluable support for help with animal experiments and colony maintenance.

The authors express gratitude to Frisén lab members – Helena Lönnqvist for endless help throughout this work and Sarantis Giatrellis for support with flow cytometry experiments.

Work was supported by EMBO Post-doctoral Fellowship (ALTF 337-2021) and MSCA Post-doctoral Fellowship (Grant agreement number 101066953) to ML, The Ming Wai Lau Centre for Reparative Medicine (MWLC) Seed Grant to JF.

