## Supplementary material for "Reconstitution of the Spinal Cord Injury Microenvironment in Adult Neural Stem Cell-Derived Organoids"

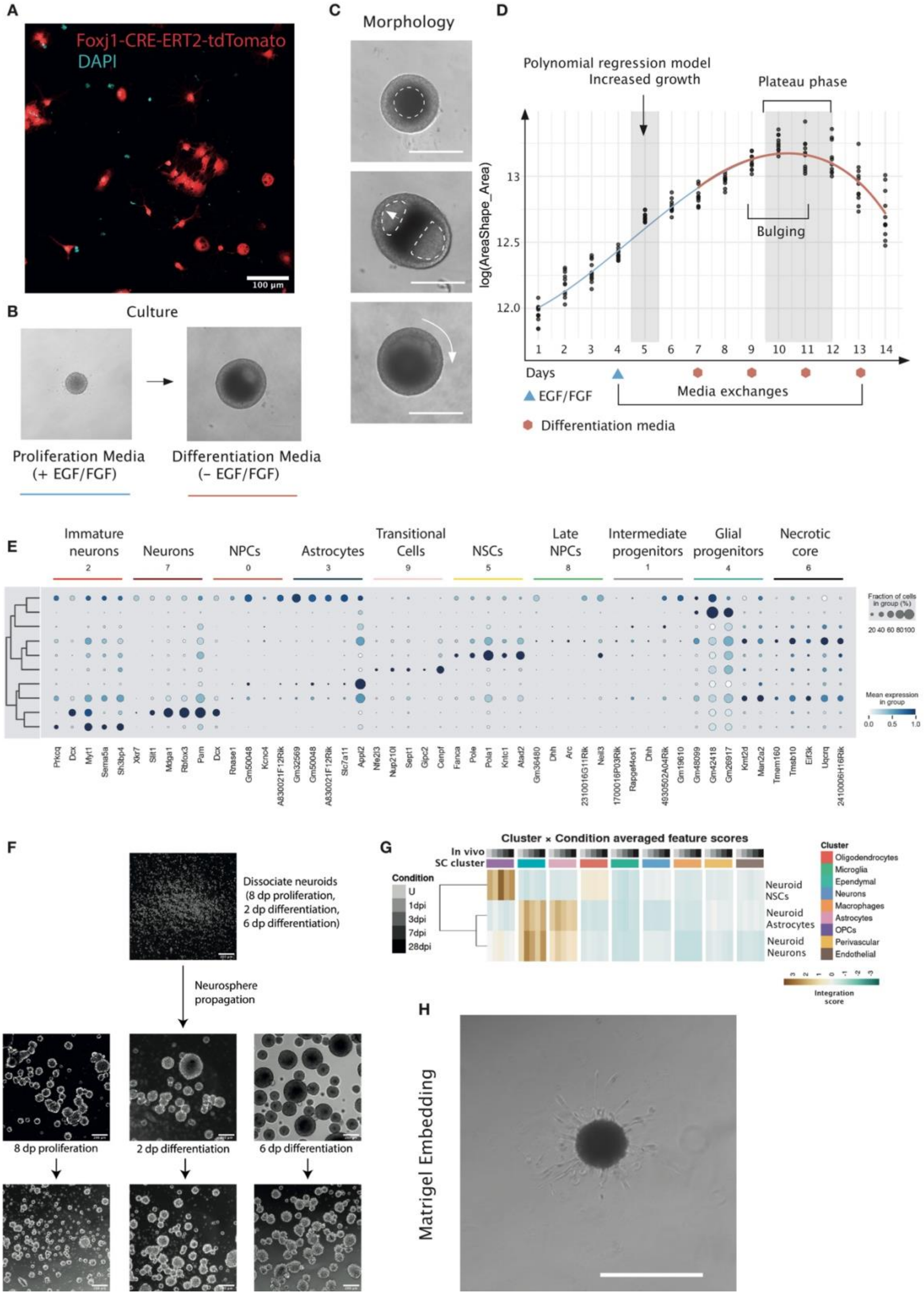

**Figure S1 | Neuroid Formation and Characterisation. (A)** A confocal image of Foxj1-CreERT2-tdTomato lineage traced spinal cord neural stem cells cultured in in vitro neurosphere cultures stained for tdTomato (in red) and nuclei (DAPI in cyan). Scale bar = 100 µm. **(B)** Brightfield images showing neuroid growth under proliferation conditions (EGF/FGF) followed by differentiation induction after growth factor withdrawal. Scale bar = 500 µm. **(C)** Brightfield images showing morphological changes of neuroids during culture including early spherical formation, transitional morphology and later stage compact neuroid structure. Dashed lines indicate neuroid phenotypical structures. Scale bar = 200 µm. **(D)** Polynomial regression model describing neuroid growth dynamics measured as log(area_shape/area) across culture days. Increased growth phase and plateau phase are indicated. Blue triangles represent EGF/FGF supplementation and red hexagons indicate differentiation media changes. **(E)** Dot plot showing expression of selected marker genes across identified neuroid clusters including immature neurons, neurons, neural progenitor cells (NPCs), astrocytes, transitional cells, neural stem cells (NSCs), late NPCs, intermediate progenitors, glial progenitors and necrotic core cells. Dot size represents fraction of expressing cells and colour intensity represents mean expression. **(F)** Brightfield images showing neuroid dissociation and neurosphere propagation after different neuroid differentiation stages (8 days post proliferation induction, 2 days and 6 days post differentiation induction). Scale bar = 200 µm. **(G)** Heatmap of neuroid neural stem cell (NSC), astrocyte and neuron cell integration scoring based on chromatin accessibility with in vivo spinal cord cells (U – uninjured, 1, 3, 7, 28 days post injury (dpi). **(H)** Brightfield image showing neuroid embedding in Matrigel followed by radial cell outgrowth. Scale bar = 200 µm.

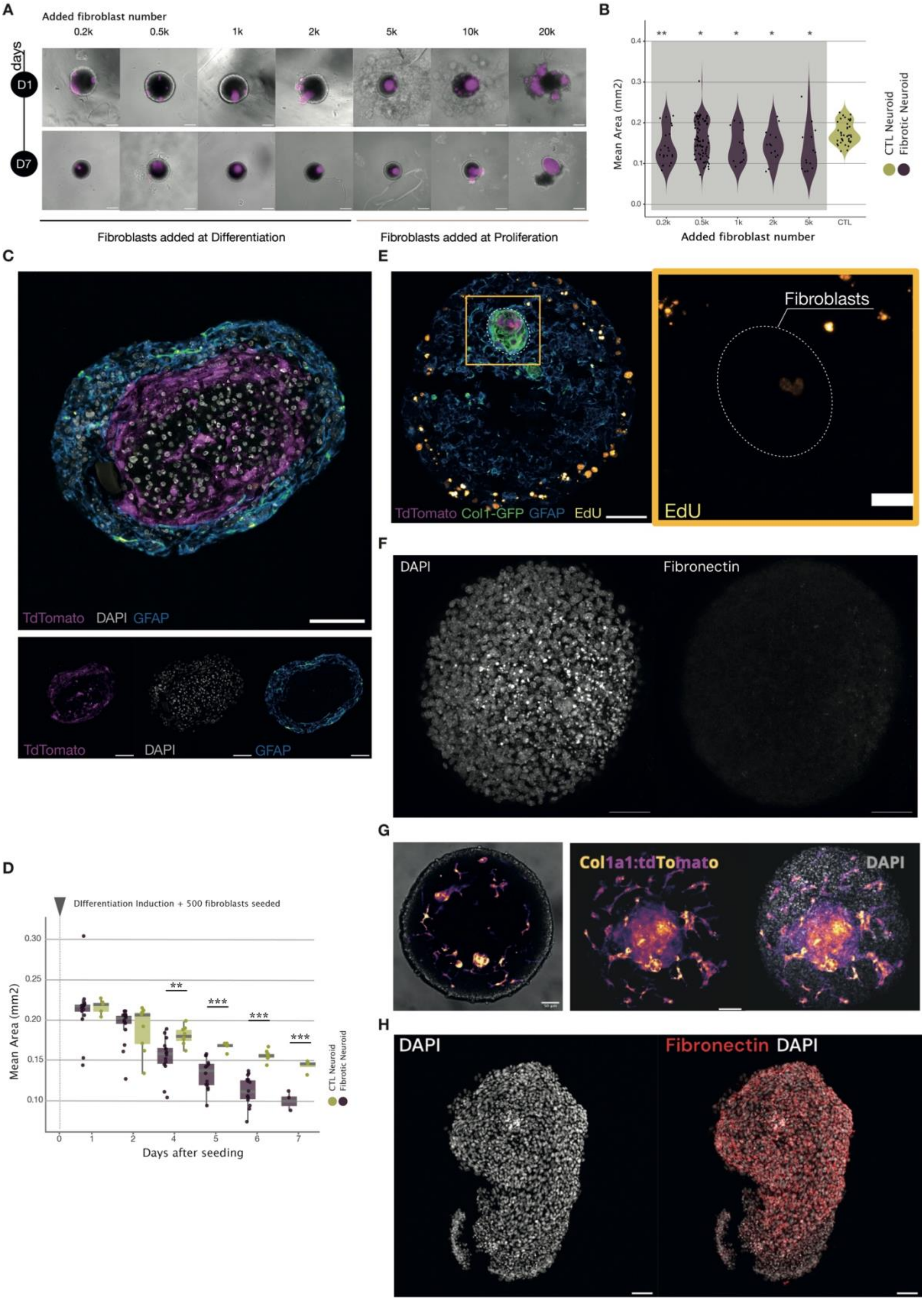

A
Added fibroblast number
0.2k
0.5k
1k
2k
5k
10k
20k
days
D1
D7
Fibroblasts added at Differentiation
Fibroblasts added at Proliferation
B
Mean Area (mm2)
Added fibroblast number
CTL Neuroid
Fibrotic Neuroid
C
TdTomato DAPI GFAP
D
Differentiation Induction + 500 fibroblasts seeded
Mean Area (mm2)
Days after seeding
CTL Neuroid
Fibrotic Neuroid
E
Fibroblasts
TdTomato Col1-GFP GFAP EdU
EdU
F
DAPI
Fibronectin
G
Col1a1:tdTomato
DAPI
H
DAPI
Fibronectin DAPI

**Figure S2 | Fibroblast Integration and Extracellular Matrix Deposition in Fibrotic Neuroids.** **(A)** Brightfield images showing effects of increasing numbers of seeded fibroblasts (0.2k–20k cells) on neuroid morphology at day 1 and day 7 after seeding. Fibroblasts are labelled in magenta. Scale bar = 200 µm. **(B)** Violin plot showing mean neuroid area across different fibroblast seeding numbers compared to control neuroids (CTL). Statistical comparisons between each fibrotic condition and the CTL group were performed using an unpaired Student's t-test. Significance levels are denoted as: $*p < 0.05$, $**p < 0.005$. Sample size: n = 22 to 80 from two technical replicates across four independent experiments. **(C)** Confocal images of fibrotic neuroid section stained for lineage traced fibroblasts (tdTomato in magenta), astrocytes (GFAP in cyan) and nuclei (DAPI in grey). Bottom panels show separated channels. Scale bar = 50 µm. **(D)** Box plots showing quantification of neuroid area over time after 500 fibroblast seeding following differentiation induction. Individual data points correspond to biological replicates (n = 6 –15 neuroids per time point). Statistical differences between time points were assessed using an unpaired Student's t-test. Significance levels are indicated as: $**p < 0.01$, $***p < 0.01$. **(E)** Confocal image of fibrotic neuroid showing proliferating fibroblasts detected by EdU incorporation (yellow), fibroblast lineage tracing (tdTomato in magenta), Col1a1-GFP reporter (green) and astrocytes (GFAP in cyan). Scale bar = 50µm Yellow box indicates zoomed region highlighting fibroblasts that have proliferated. Scale bar = 20µm **(F)** Confocal images showing control neuroids stained for nuclei (DAPI) and extracellular matrix protein fibronectin showing no fibronectin deposition in the absence of fibroblasts. Scale bar = 50µm. **(G)** Confocal images showing neuroids populated with lineage traced Col1a1-CreERT2-tdTomato (in yellow-magenta) skeletal muscle fibroblasts. **(H)** Confocal images showing structural neuroid destruction by using mouse embryonic fibroblast cell line, that still deposit fibronectin (in red), stained nuclei (DAPI in grey). Scale bar = 50µm.

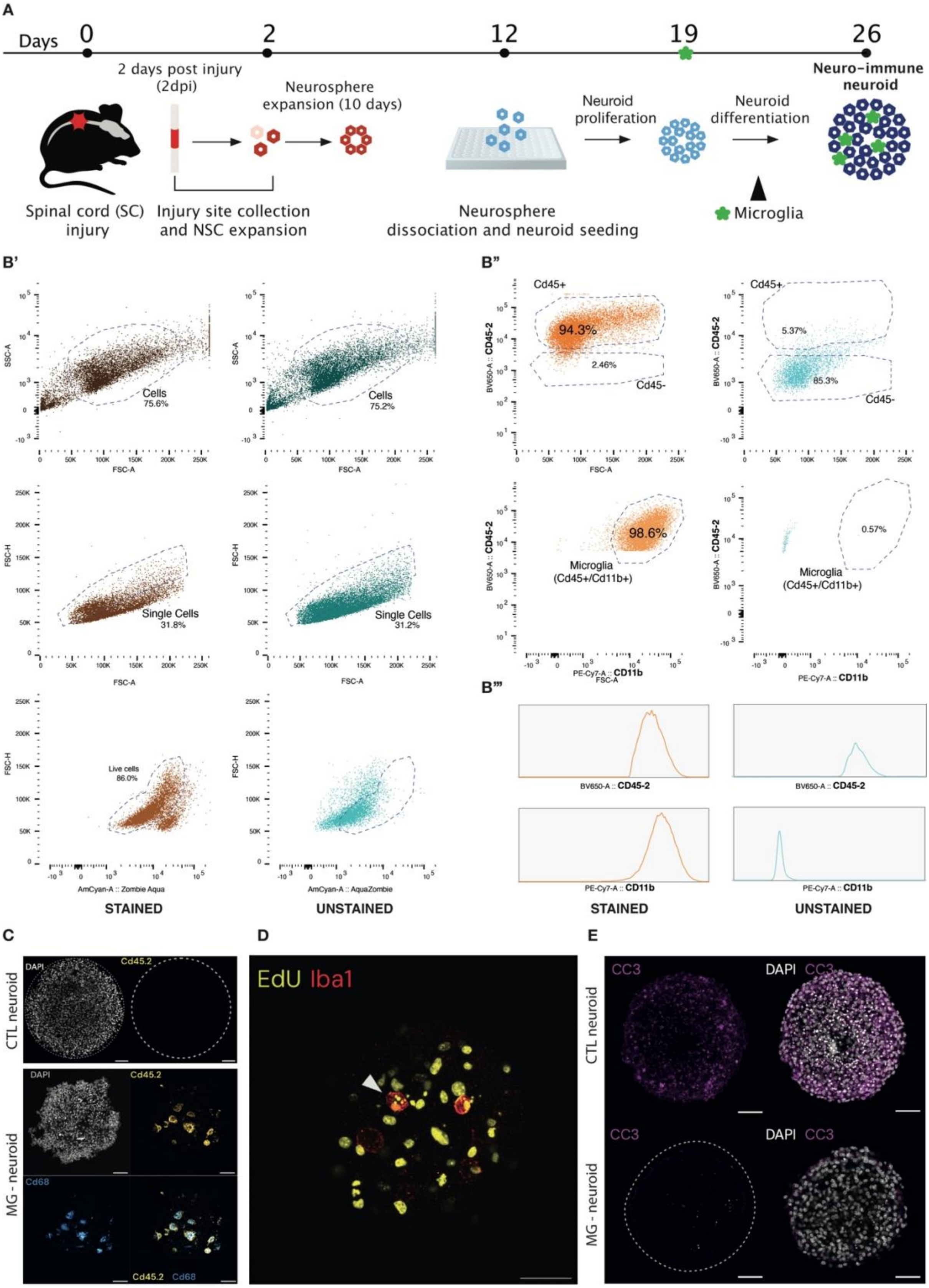

A
Days 0 2 12 19 26
2 days post injury (2dpi)
Neurosphere expansion (10 days)
Neuroid proliferation
Neuroid differentiation
Neuro-immune neuroid
Spinal cord (SC) injury
Injury site collection and NSC expansion
Neurosphere dissociation and neuroid seeding
Microglia
B'
Cells 75.6%
Cells 75.2%
Single Cells 31.8%
Single Cells 31.2%
Live cells 86.0%
SSC-A
FSC-A
FSC-H
AmCyan-A :: Zombie Aqua
AmCyan-A :: AquaZombie
STAINED
UNSTAINED
B"
Cd45+
94.3%
2.46%
Cd45-
5.37%
85.3%
BV650-A :: CD45-2
98.6%
0.57%
Microglia (Cd45+/Cd11b+)
PE-Cy7-A :: CD11b
B'''
BV650-A :: CD45-2
PE-Cy7-A :: CD11b
STAINED
UNSTAINED
C
CTL neuroid
MG - neuroid
DAPI
Cd45.2
Cd68
Cd45.2 Cd68
D
EdU Iba1
E
CC3
DAPI CC3
CTL neuroid
MG - neuroid

**Figure S3 | Microglia Isolation and Validation for Neuroid Incorporation. (A)** Schematics depicting experimental workflow for isolation of adult microglia and incorporation into neuroids. **(B')** Flow cytometry gating strategy identifying total cells, single cells and live cells in stained and unstained cultured microglial preparations. **(B'')** Top plots - flow cytometry plots showing CD45 positive microglial population and negligent staining in unstained samples. Bottom plots - CD45/Cd11b double positive microglial population and no detected staining in unstained samples. **(B''')** Histograms showing expression of CD45 and CD11b in stained and unstained samples confirming microglial identity. **(C)** Confocal images of control neuroids and MG-neuroids stained for CD45. Microglial markers CD68 and CD45 are shown in MG-neuroids. Scale bar = 50µm. **(D)** Confocal image showing engulfed pyknotic EdU+ (yellow) nuclei in infiltrated microglia labelled by Iba1 (red), demonstrating their phagocytic properties. Scale bar = 50µm. **(E)** Confocal images of neuroids stained for cleaved caspase-3 (CC3, in magenta) indicating apoptotic cells in control and MG-neuroids. Nuclei are shown with DAPI (in grey). Scale bar = 50µm.

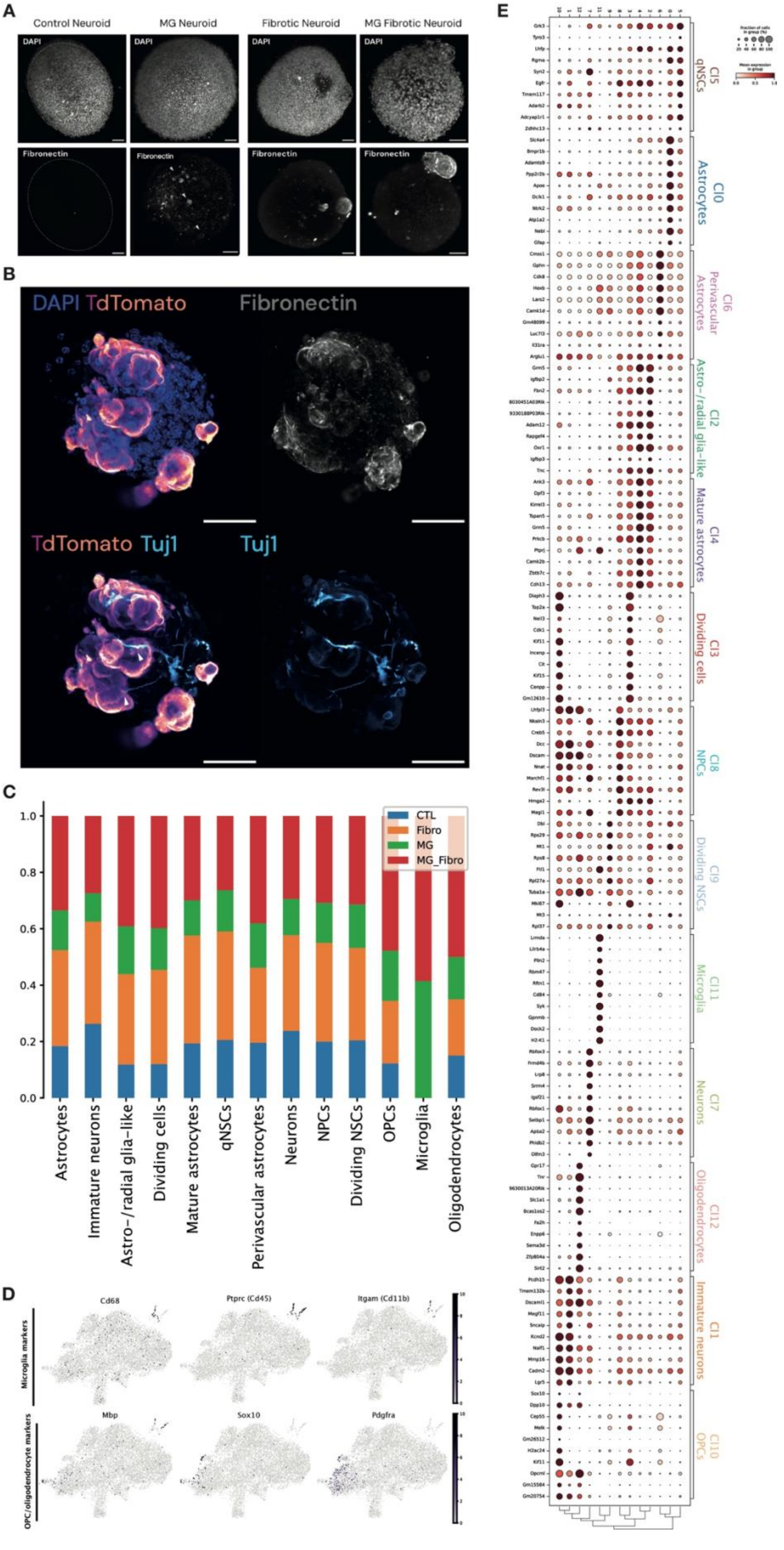
A
Control Neuroid
MG Neuroid
Fibrotic Neuroid
MG Fibrotic Neuroid
DAPI
Fibronectin
B
DAPI TdTomato
Fibronectin
TdTomato Tuj1
Tuj1
C
CTL
Fibro
MG
MG_Fibro
1.0
0.8
0.6
0.4
0.2
0.0
Astrocytes
Immature neurons
Astro-/radial glia-like
Dividing cells
Mature astrocytes
qNSCs
Perivascular astrocytes
Neurons
NPCs
Dividing NSCs
OPCs
Microglia
Oligodendrocytes
D
Microglia markers
Cd68
Ptprc (Cd45)
Itgam (Cd11b)
OPC/oligodendrocyte markers
Mbp
Sox10
Pdgfra
E
Cl5 qNSCs
Cl0 Astrocytes
Cl6 Perivascular Astrocytes
Cl2 Astro-/radial glia-like
Cl4 Mature astrocytes
Cl3 Dividing cells
Cl8 NPCs
Cl9 Dividing NSCs
Cl11 Microglia
Cl7 Neurons
Cl12 Oligodendrocytes
Cl1 Immature neurons
Cl10 OPCs

**Figure S4 | Integrated Multi-Condition Neuroid Dataset Characterisation. (A)** Confocal images of neuroids from different experimental conditions including control neuroids (CTL), microglia-seeded neuroids (MG neuroids), fibrotic neuroids (Fibrotic neuroids) and neuroids containing both microglia and fibroblasts (MG-Fibrotic neuroids). Top panels show nuclei staining (DAPI in grey). Bottom panels show fibronectin staining highlighting extracellular matrix deposition within migroclia containing neuroids and neuroid with fibrotic conditions. Dashed circle outlines the control neuroid boundary. Scale bar = 50 µm. **(B)** Confocal images of fibrotic neuroids stained for nuclei (DAPI in blue), lineage traced fibroblasts (Col1a1-CreERT2-tdTomato in magenta) and neuronal marker β-III tubulin (Tuj1 antibody, in cyan). Right panels show separated channels for fibronectin and Tuj1 staining. Scale bar = 50 µm. **(C)** Stacked barplots showing cluster composition across neuroid conditions (CTL, Fibro, MG and MG-Fibro). Each bar represents the fraction of cells assigned to annotated clusters including astrocytes, immature neurons, astro-/radial glia-like cells, dividing cells, mature astrocytes, qNSCs, perivascular astrocytes, neurons, NPCs, dividing NSCs, OPCs, microglia and oligodendrocytes. **(D)** Feature plots visualised in UMAP space showing expression of selected marker genes for microglia (Cd68, Ptprc (Cd45), Itgam (Cd11b)) and oligodendrocyte lineage cells (Mbp, Sox10, Pdgfra). **(E)** Dot plot showing expression of canonical marker genes across identified Leiden clusters. Dot size represents the fraction of cells expressing the gene and colour intensity represents mean expression level. Annotated clusters include qNSCs, astrocytes, perivascular astrocytes, astro-/radial glia-like cells, mature astrocytes, dividing cells, NPCs, dividing NSCs, microglia, neurons, oligodendrocytes, immature neurons and OPCs.

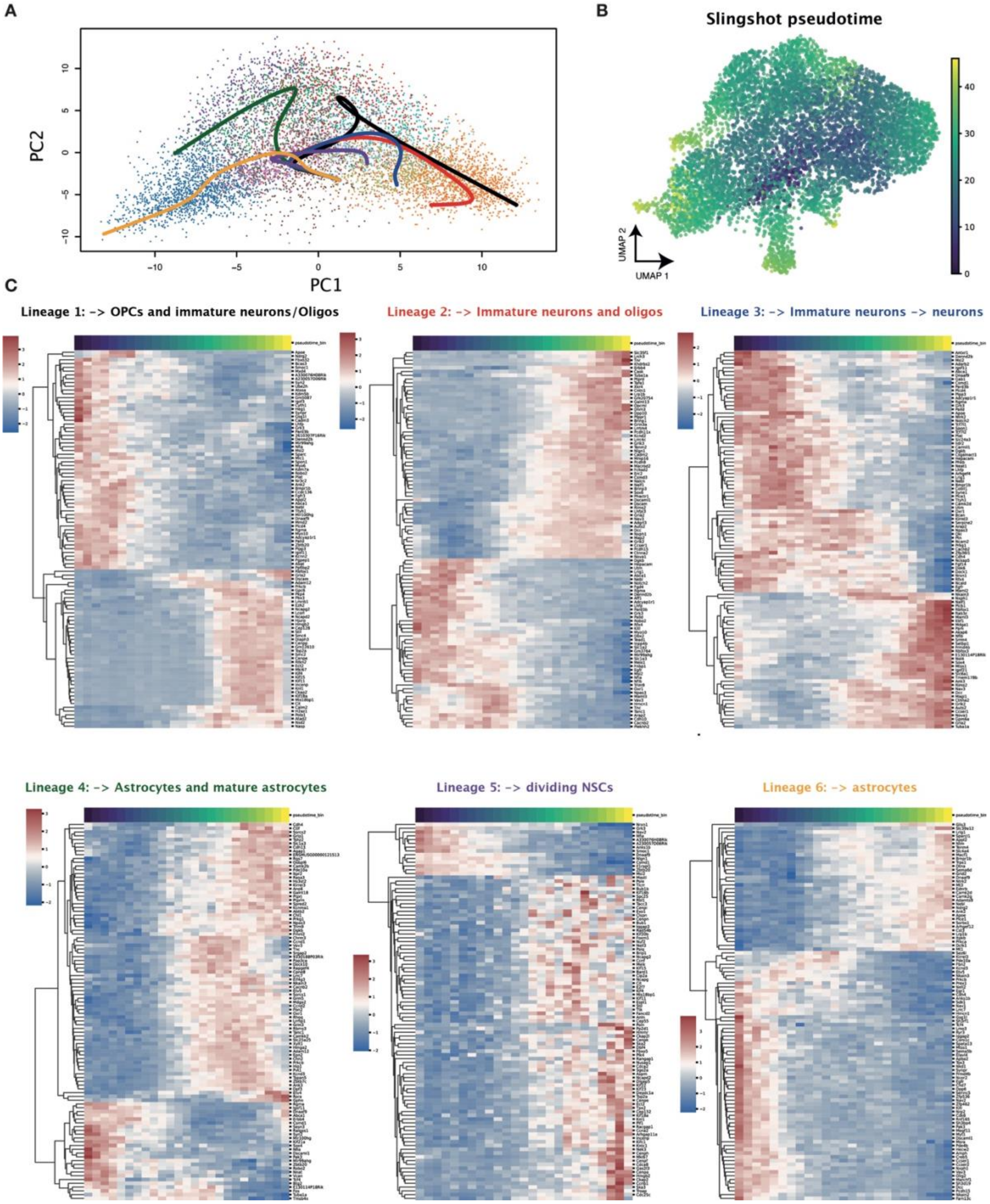


**Figure S5 | Neural Stem Cell Lineage Reconstruction and Pseudotime Gene Expression Dynamics. (A)** Slingshot-inferred lineage trajectories fitted on PCA embeddings showing predicted differentiation paths emerging from quiescent neural stem cell cluster. **(B)** Slingshot pseudotime visualised in UMAP space showing progression of cells along differentiation trajectories. **(C)** Heatmaps showing gene expression dynamics along pseudotime for each identified lineage. Colour scale represents relative gene expression across pseudotime progression.

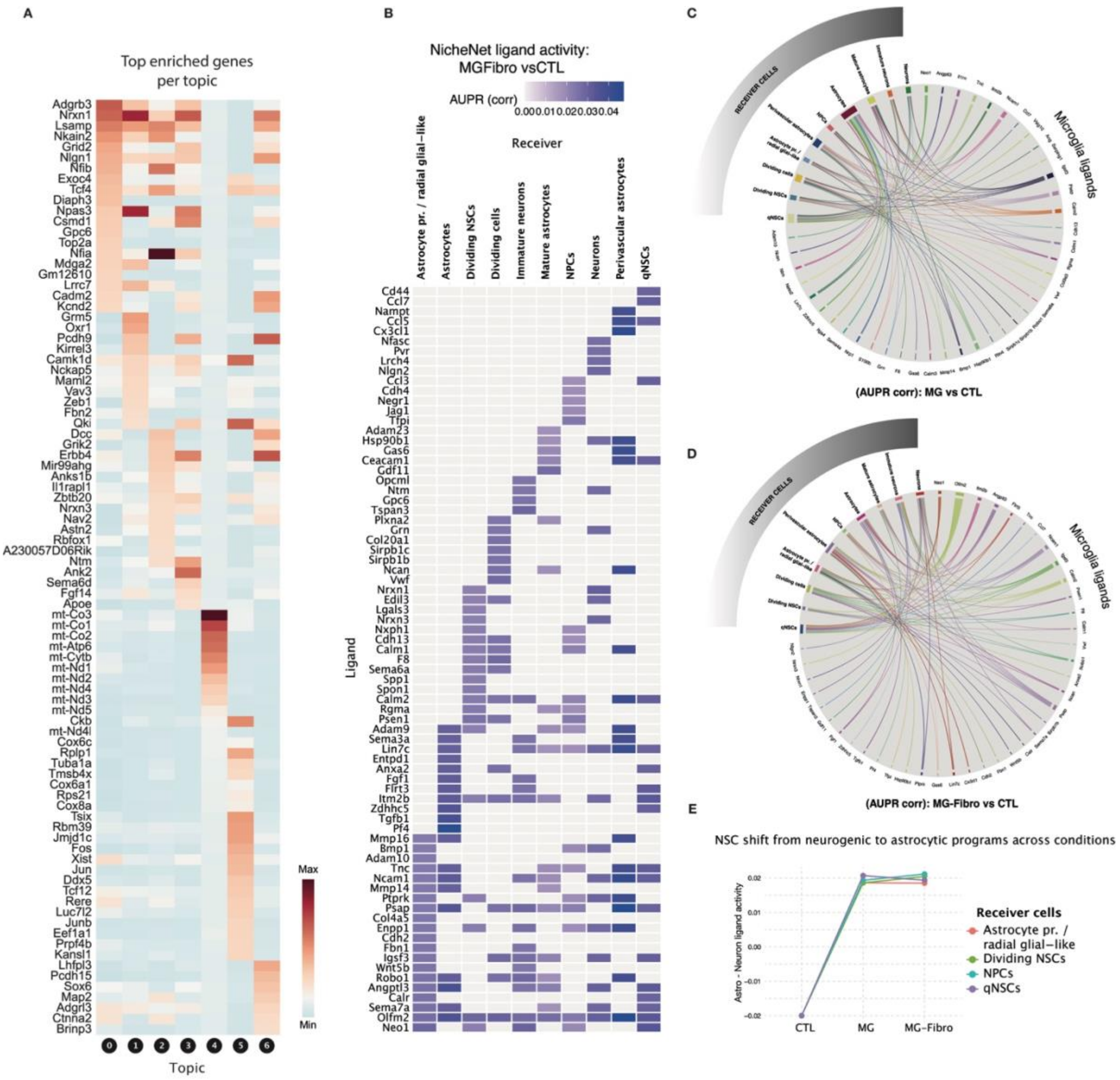


**Figure S6 | Topic-Associated Gene Programs and Microglial Ligand Signalling. (A)** Heatmap showing top enriched genes associated with each topic identified by topic modelling. **(B)** Heatmap showing NicheNet predicted ligand activity for microglial ligands across neuroid receiver populations in the neuroids with microglia and fibroblasts (MG-Fibro) vs control (CTL) neuroid comparison. Ligands are shown on the Y-axis and receiver populations on the X-axis. Colour intensity represents ligand activity score (AUPR). **(C)** Circos plot visualising predicted ligand-receiver interactions between microglial ligands and neuroid cell populations in MG vs CTL comparison. **(D)** Circos plot visualising predicted ligand-receiver interactions in MG-Fibro vs CTL comparison. **(E)** Line plot showing neural stem cell program shift from neurogenic toward astrocytic transcriptional programs across neuroid conditions (CTL, MG and MG-Fibro). Receiver populations include astrocyte progenitors / radial glia-like cells, dividing NSCs, NPCs and qNSCs.

**Supplementary Table 1: List of antibodies and dyes**

| Primary antibodies | Company | Cat. No. | Dillution | Species | Clonality |
|---|---|---|---|---|---|
| anti-Iba1 | Wako | 019-19741 | 1:500 | Rabbit | Polyclonal |
| anti-Cd68 | Abcam | ab53444 | 1:200 | Rat | Monoclonal |
| anti-Cd45.2 | eBioscience | AB_467261 | 1:200 | Mouse | Monoclonal |
| anti-GFAP | Dako | Z0334 | 1:500 | Rabbit | Polyclonal |
| anti-βIII-tubulin/TUJ1 | BioLegend | AB_2313773 | 1:200 | Mouse | Monoclonal |
| anti-Ki67 | eBioscience | 14-5698-82 | 1:500 | Rat | Monoclonal |
| anti-Fibronectin | Milipore | AB2033 | 1:200 | Rabbit | Polyclonal |
| anti-Fibronectin | BioRad | VPA00045 | 1:200 | Sheep | Polyclonal |
| anti-Collagen I | Abcam | ab21286 | 1:100 | Rabbit | Polyclonal |
| anti-Caspase 3 | Invitrogen | AB_2745900 | 1:200 | Rabbit | Monoclonal |
| anti-RFP | Rockland | 600-901-379 | 1:1000 | Chicken | Polyclonal |
| anti-GFP | Rockland | 600-101-215M | 1:1000 | Goat | Monoclonal |

All seconady antibodies were raised in donkey:

| Secondary antibodies | Company | Cat. No. | Dillution |
|---|---|---|---|
| Alexa Fluor® 488 AffiniPure® Donkey Anti-Goat IgG (H+L) | Jackson ImmunoResearch | 705-545-147 | 1:500 |
| Alexa Fluor® 488 AffiniPure® Donkey Anti-Rat IgG (H+L) | Jackson ImmunoResearch | 705-545-150 | 1:500 |
| Alexa Fluor® 488 AffiniPure® Donkey Anti-Sheep IgG (H+L) | Jackson ImmunoResearch | 713-545-147 | 1:500 |
| Cy™3 AffiniPure® Donkey Anti-Rabbit IgG (H+L) | Jackson ImmunoResearch | 711-165-152 | 1:500 |
| Cy™3 AffiniPure® Donkey Anti-Chicken IgY (IgG) (H+L) | Jackson ImmunoResearch | 703-165-155 | 1:500 |
| Cy™3 AffiniPure® Donkey Anti-Mouse IgG (H+L) | Jackson ImmunoResearch | 715-165-151 | 1:500 |
| Alexa Fluor® 647 AffiniPure® Donkey Anti-Mouse IgG (H+L) | Jackson ImmunoResearch | 715-605-150 | 1:500 |
| Alexa Fluor® 647 AffiniPure® Donkey Anti-Rabbit IgG (H+L) | Jackson ImmunoResearch | 711-605-152 | 1:500 |
| Alexa Fluor® 647 AffiniPure® Donkey Anti-Rat IgG (H+L) | Jackson ImmunoResearch | 712-605-150 | 1:500 |

| Dyes | Company | Cat. No. | Dillution |
|---|---|---|---|
| DAPI | Thermo Scientific | 62248 | 1:500 |
